# Nuclear mRNA metabolism drives selective basket assembly on a subset of nuclear pores in budding yeast

**DOI:** 10.1101/2021.11.07.467636

**Authors:** Pierre Bensidoun, Taylor Reiter, Ben Montpetit, Daniel Zenklusen, Marlene Oeffinger

## Abstract

To determine which transcripts should reach the cytoplasm for translation, eukaryotic cells have established mechanisms to regulate selective mRNA export through the nuclear pore complex (NPC). The nuclear basket, a substructure of the NPC protruding into the nucleoplasm, is thought to function as a stable platform where mRNA-protein complexes (mRNPs) are rearranged and undergo quality control (QC) prior to export, ensuring that only mature mRNAs reach the cytoplasm. Here, we use proteomic, genetic, live-cell, and single-molecule resolution microscopy approaches in budding yeast to demonstrate that baskets assemble only on a subset of NPCs and that basket formation is dependent on RNA polymerase II (Pol II) transcription and subsequent mRNP processing. Specifically, we observe that the cleavage and polyadenylation machinery, the poly(A)-binding protein Pab1, and pre-mRNA-leakage factor Pml39 are required for basket assembly. We further show that while all nuclear pores can bind Mlp1, baskets assemble only on a subset of nucleoplasmic NPCs, and these basket-containing pores associate a distinct protein and RNA interactome. Taken together, our data points towards nuclear pore heterogeneity and an RNA-dependent mechanism for functionalization of nuclear pores in budding yeast through nuclear basket assembly.

## INTRODUCTION

Exchange of macromolecules between nucleus and cytoplasm occurs through the NPC, a large multi-protein complex assembled by ~30 different proteins (called nucleoporins or nups), which forms a transport channel that spans the nuclear envelope (Alber et al., 2007; Kim et al., 2018). Transport through the NPC is mediated by transport receptors that bind their cargos and facilitate movement across the NPC by interacting with phenylalanine-glycine (FG) repeat-containing proteins (FG nups) that line the inside of the NPC’s central transport channel (Lin et al., 2019). Cargo access to and release from the NPC is further modulated by asymmetrically distributed subcomplexes of the NPC. On the nuclear face, this is accomplished by a large basket-like structure protruding ~100nm into the nucleoplasm, termed the nuclear basket (Bensidoun et al., 2021).

The basket’s main scaffold is assembled by the filamentous protein TPR (Translocated Promoter Region protein) in humans and the two paralogues Mlp1 and Mlp2 (myosin-like protein) in budding yeast (Frosst et al., 2002; Galy et al., 2004; Zhao et al., 2004). Mlp1/Mlp2 and TPR are large proteins with predicted coiled-coil regions spanning approximately two-thirds of the proteins, with a long intrinsically disordered C-terminal domain (IDD). The coiled-coil regions are thought to form the spokes of the basket anchoring it at the NPC, whereas the C-terminus forms the top of the basket, the so-called distal ring. Yet unlike for the central framework of the NPC, high resolution structures of the nuclear basket have not yet been obtained, nor has the exact stoichiometry of these proteins at individual NPCs been determined (Alber et al., 2007; Kim et al., 2018; Mi et al., 2015; Rajoo et al., 2018).

The nuclear basket has been shown to contribute to a range of nuclear activities, including modulation of DNA topology, DNA repair, epigenetic regulation, and anchoring of genes at the nuclear periphery; however, the main role of the basket in gene regulation is thought to be mediating the access of messenger ribonucleoproteins (mRNPs) to the NPC (Brickner et al., 2004; Pascual-Garcia et al., 2019; Raices et al., 2012; Taddei et al., 2006). Early studies investigating the large Balbiani Ring mRNPs expressed in salivary glands of *Chironomus tentans* showed that mRNPs associate with the top of the basket prior to decompacting and with the 5’ end of the mRNA entering the NPC first, leading to a model where association with the basket acts as a rate limiting step of RNA transport and a site of mRNP reorganization (Björk et al., 2015). Various studies have since demonstrated mRNP reorganization involving basket-associated proteins, and single molecule studies further confirmed the involvement of the basket in a rate limiting step at the NPC as part of mRNP export (Björk et al., 2017; Bretes et al., 2014; Grünwald et al., 2010; Iglesias et al., 2010; Mor et al., 2010; Saroufim et al., 2015).

Surprisingly, however, Mlp1/2 and TPR are not required for mRNA export per se, as their loss only leads to a mild or partial export defect, respectively, indicating that the basket may only facilitate export of some transcripts, or has additional functions (Green et al., 2003; Powrie et al., 2011; Saroufim et al., 2015; Soucek et al., 2016; Zander et al., 2016; Aksenova et al., 2020; Lee et al., 2020; Bensidoun et al., 2021). One such proposed function is that of establishing a quality control platform that ensures that only mature mRNPs are exported to the cytoplasm, as deletion of Mlp1 was shown to result in the leakage of intron-containing mRNAs to the cytoplasm, suggesting that the basket can selectively grant NPC access to spliced but not pre-mRNAs (Bonnet et al., 2015; Galy et al., 2004). While the mechanism of this process is not yet fully understood, it might involve_RNA-binding proteins (RBPs) associating with mRNPs that serve as signals for export or retention, possibly modulating the ability of mRNPs to interact with baskets (Hackmann et al., 2014). Consistent with such a model, in yeast, various RBPs showing pre-mRNA leakage phenotypes when mutated and either associating with pre-mRNPs or Mlp1/baskets have been identified, including Gbp1, Hrb1, the Pre-mRNA Leakage proteins Pml1 and Pml39, as well as the nuclear envelope protein Esc1 and the basket protein Nup60 (Palancade et al., 2005; Bonnet et al., 2015; Dziembowski et al., 2004; Hackmann et al., 2014; Lewis et al., 2007). Moreover, different RBPs required for mRNA export were shown to interact with Mlp1, including the nuclear poly(A)-binding protein (PABP) Nab2, which directly interacts with the C-terminus of Mlp1, further pointing towards the basket as a site of late mRNA maturation and/or initial contact point between mRNPs and NPCs to regulate export (Green et al., 2003; Saroufim et al., 2015; Soucek et al., 2016; Zander et al., 2016).

While one might expect of such a platform to be stably anchored to the rest of the NPC, fluorescence recovery after photobleaching (FRAP) experiments have shown that exchange of Mlp1 at NPCs is faster when compared to other nups (Niepel et al., 2005; Niño et al., 2016). Moreover, Mlp1 and Mlp2 were shown to dissociate from NPCs during heat-shock at 42°C and assemble into intra-nuclear granules together with mRNA maturation factors, including Nab2 and Yra1, and poly(A) mRNAs (Carmody et al., 2010; Zander et al., 2016). The reasons for the dynamic association of Mlp1/2 with the pore are still unknown, and the mechanisms leading to the formation of Mlp1 granules as well as their function are poorly understood. Moreover, in *S. cerevisiae,* NPCs adjacent to the nucleolus, which occupies about a third of the nuclear volume, are devoid of baskets (Miné-Hattab et al., 2019; Galy et al., 2004; Niepel et al., 2013). Yet how cells establish these basket-less pores, and whether they represent specialized NPCs with functions differing from nucleoplasmic, basket-containing pores is not known.

Here, we show that, unlike previously thought, basket-less pores are not a specialized state of NPCs. Rather, our data demonstrate that basket-less pores are the default assembly state formed throughout the nuclear envelope and baskets assemble dynamically onto a subset of nucleoplasmic pores in an mRNA-dependent manner. Specifically, inhibition of RNA polymerase II transcription results in the abrogation of basket assembly at nucleoplasmic NPCs, and interference with 3’ end processing and polyadenylation leads to the loss of nuclear baskets, linking specific steps of mRNA maturation to basket assembly. Moreover, these disruptions resulted either in the formation of nuclear Mlp1-containing granules or the re-location of baskets to the nucleolar periphery, the latter inducing a concomitant internalization of the nucleolus away from the periphery, suggesting an incompatibility between NPC basket assembly and the nucleolus. Proteomic, microscopy and RNA sequencing experiments further show that basket-containing pores assemble a unique protein and RNA interactome and that, while both types of pores associate with mRNAs and are likely export competent, Mlp1 preferentially associates with longer mRNAs, but not short and/or intron-containing mRNAs. Taken together, our data identifies an RNA-dependent mechanism for functionalization of nuclear pores in budding yeast through nuclear basket assembly.

## RESULTS

### Mlp1 can traverse the nucleolus and form rare ectopic baskets at the nucleolar periphery

While nuclear pores are evenly distributed along the nuclear periphery, not all parts of the NPC follow this pattern of distribution. In *S. cerevisiae,* the nuclear basket proteins Mlp1 and Mlp2 are excluded from nuclear pores next to the nucleolus, a crescent-shaped membrane-less compartment that is positioned adjacent to the nuclear membrane and is the site of ribosome biogenesis. Nucleoli, however, change with stress, cellular aging, cell cycle stage as well as overall metabolic activity, which can cause significant variability in nucleolar size and shape between cells in non-synchronous cultures (Neumüller et al., 2013; Sirri et al., 2008), suggesting that the number of nuclear pores that are bound by Mlp1/2 may also vary. While Mlp1 and Mlp2 are both considered part of the basket scaffold, Mlp2 localization at NPCs requires Mlp1, but not vice versa (Palancade et al., 2005); we therefore focussed our study on Mlp1. Measuring the distribution of Mlp1-GFP to the nucleolar marker Gar1-tdTomato along the circumference of the nuclear envelop, we observed a negative correlation between the nucleolar and Mlp1-occupied territories (Figures 1A, B). This suggests that variations in nucleolar size negatively affect the number of pores available to bind Mlp1 and, moreover, that the number of basket-containing pores is not fixed but instead may vary from cell to cell and over time.

**Figure 1.**
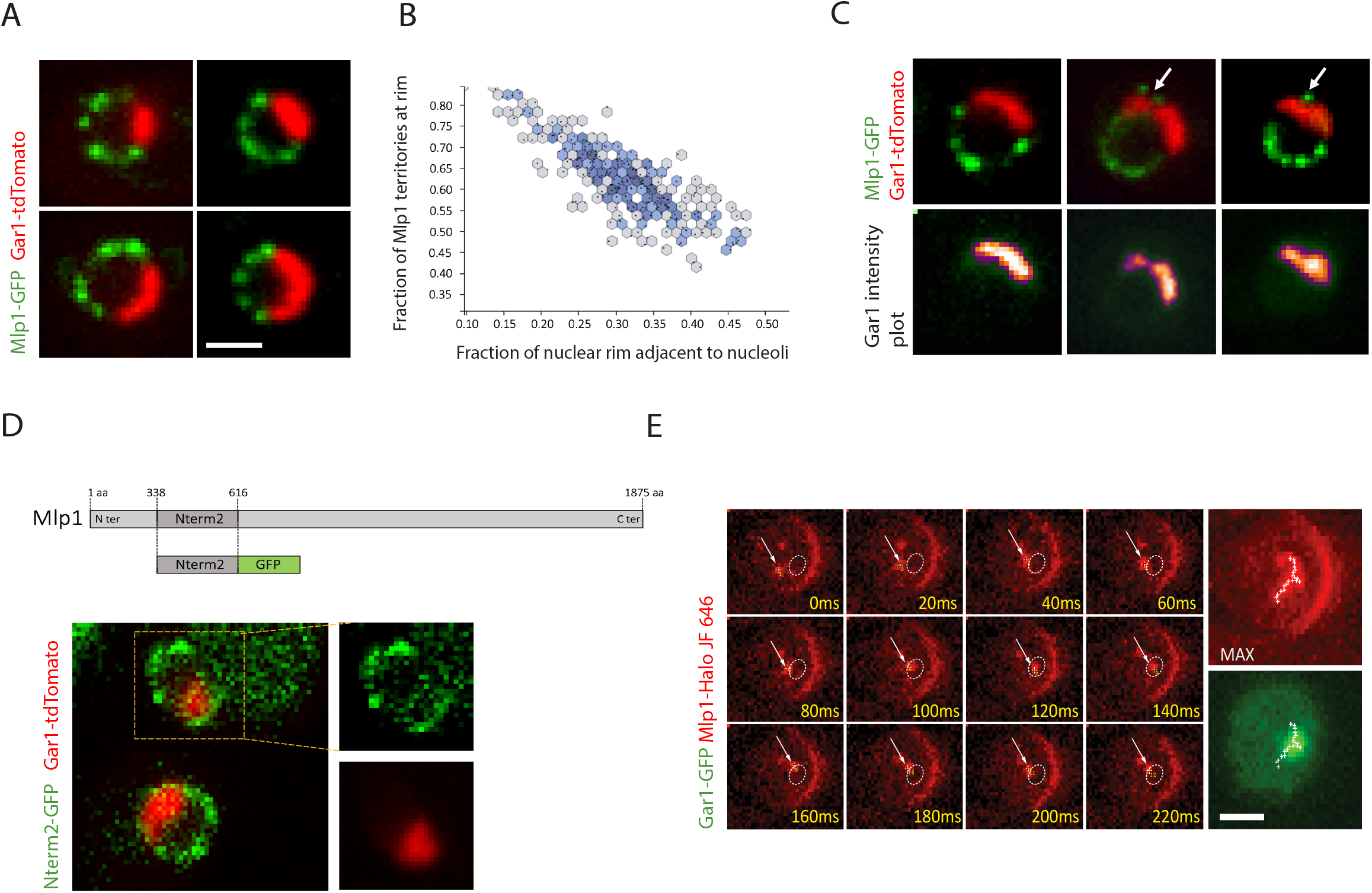
Mlp1 can access nucleolar pores and assemble baskets in the nucleolus. **A.** Fluorescent microscopy images showing Mlp1-GFP (green) and Gar1-tdTomato (red) distribution in non-synchronized cultures. **B.** Quantification of the percentage of the nuclear periphery occupied by Mlp1-GFP versus the percentage occupied by the nucleolus (n=200 cells) **C.** Mlp1-GFP can assemble an ectopic basket at the nuclear periphery adjacent to nucleoli (white arrows). Ectopic baskets coincide with a loss of Gar1-tdTomato signal (Gar1 intensity plot). **D**. Cartoon of the Mlp1 fragment containing an NPC binding domain (Nterm2) spanning a 278 amino acids region. The Nterm2-GFP fragment can bind the periphery including the nucleolar area in *Δ mlp1/2.* Nucleolus marked with Gar1-tdTomato strain (red). **E.** Live-cell tracking of single Mlp1-Halo JF646 molecules. Individual frames acquired at 20ms intervals are shown. White arrows show Mlp1-Halo JF646 and the dashed circle presents the nucleolar area. MAX shows the maximum intensity projection of all frames of the movie with the track highlighted with white crosses. Mlp1-Halo JF646 is shown in red, nucleolus (Gar1-GFP) in green. (Scale bar = 2μm).

Although Mlp1 is generally excluded from the nucleolus, in ~1% of cells, we observed an Mlp1-GFP focus along the nucleolar-occupied territory, here referred to as an ectopic basket (Figure 1C). The presence of an ectopic basket correlated with an invagination in the nucleolar signal surrounding the Mlp1 focus, indicating that baskets can assemble along the nucleolar periphery but that these territories are mutually exclusive and cannot co-occur. Ectopic baskets may be observed when Mlp1 assembles onto a nucleolar NPC or a basket-containing nuclear pore is isolated by the process of nucleolar expansion. To determine if Mlp1 can associate with nucleolar NPCs, we investigated whether an N-terminal fragment of Mlp1 (N-terminal fragment 2), previously shown to be sufficient to associate with NPCs, was able to bind nuclear pores along the nucleolar periphery (Niepel et al., 2013). Mlp1 N-terminal 2 region fused to GFP was ectopically expressed in an MLP1 deletion background to avoid competition with endogenous Mlp1. The N-terminal 2 region of Mlp1 was able to bind NPCs both in the nucleoplasm as well as in the nucleolus (Figure 1D), suggesting that all nuclear pores along the nucleolar periphery are, in principle, competent to bind Mlp1.

A hallmark of liquid-phase separated compartments, such as nucleoli, is their ability to concentrate or to exclude specific factors depending on their biochemical properties (Feric et al., 2016; Miné-Hattab et al., 2019; Sirri et al., 2008). Hence, the nucleolus could represent a diffusion barrier for full-length Mlp1. FRAP experiments have shown that Mlp1 associates dynamically with the NPC and exists in two pools within the nucleus: an NPC-bound fraction and a free pool that in theory could also assemble baskets on NPCs in the nucleolus (Iglesias et al., 2010; Niepel et al., 2013). We therefore analyzed diffusion patterns of free Mlp1 using single-protein tracking of Halo-tagged Mlp1 and Halo-NLS bound by JF-549 in cells expressing Gar1-GFP. To observe single proteins, images were acquired using HILO illumination until most fluorophores were bleached and single molecules could be detected for tracking (Supplementary movies S1-2). While most single Mlp1 molecules were bound to the periphery and static, a nuclear diffusing fraction could also be observed for Mlp1. Free diffusing Mlp1-Halo was able to enter the nucleolus, similar to Halo-NLS, with about 30% of the tracks overlapping with the nucleolar area (Figures 1E, S1A, B). These data suggests that Mlp1 can access the nucleolus, and NPCs along the nucleolar periphery can bind Mlp1; however, formation of baskets at nucleolar pores is rare and results in a distortion of the nucleolus, suggesting that nucleoplasm-specific processes may mediate basket assembly at pores.

### Basket assembly requires mRNA production

The main differentiating feature between the nucleoplasmic and nucleolar region is the synthesis of two distinct types of RNA: RNA Polymerase I (Pol I) synthesizing ribosomal RNA (rRNA) in the nucleolus, and RNA Polymerase II (Pol II) producing mRNA in the nucleoplasm, initiating the assembly of two distinct types of RNPs. We therefore considered the possibility that basket assembly is linked to events solely occurring in the nucleoplasm and tested whether this requires processes linked to mRNA metabolism. To that end, we constructed strains in which the large subunit of either RNA Pol I (RpbA135) or RNA Pol II (Rpb2) were tagged with an Auxin Inducible Degron cassette (AID-HA), to enable their depletion upon addition of the plant hormone auxin (Figure 2A). Depletion of RNA Pol II, but not RNA Pol I, resulted in the loss of Mlp1-GFP at the nuclear periphery and its redistribution into the nucleoplasm after 120 min (Figure 2B). Mlp1-GFP relocalization from the periphery to the nuclear interior was also observed in *rpb1-1* at non-permissive temperature (37°C) after 10 minutes (Figure 2D). Moreover, in both strains, a large fraction of cells (~80%) exhibited formation of a singular nuclear Mlp1 granule (Figures 2C and S2). Similar Mlp1 granule formation has previously been observed upon heat-shock at 42°C, where Mlp1 dissociates from pores and forms intranuclear foci that contain several mRNA maturation factors, as well as upon deletion of NUP60, which is required for the anchoring of Mlp1 at the NPC (Niepel et al., 2013; Mészáros et al., 2015). These observations link basket formation, which is restricted to the nucleoplasm, to RNA Pol II activity and hence nuclear mRNA metabolism. However, these experiments do not discriminate whether loss of transcriptional activity per se or the consequential absence of mRNA maturation and/or mRNA export is required for NPCs to assemble baskets.

**Figure 2.**
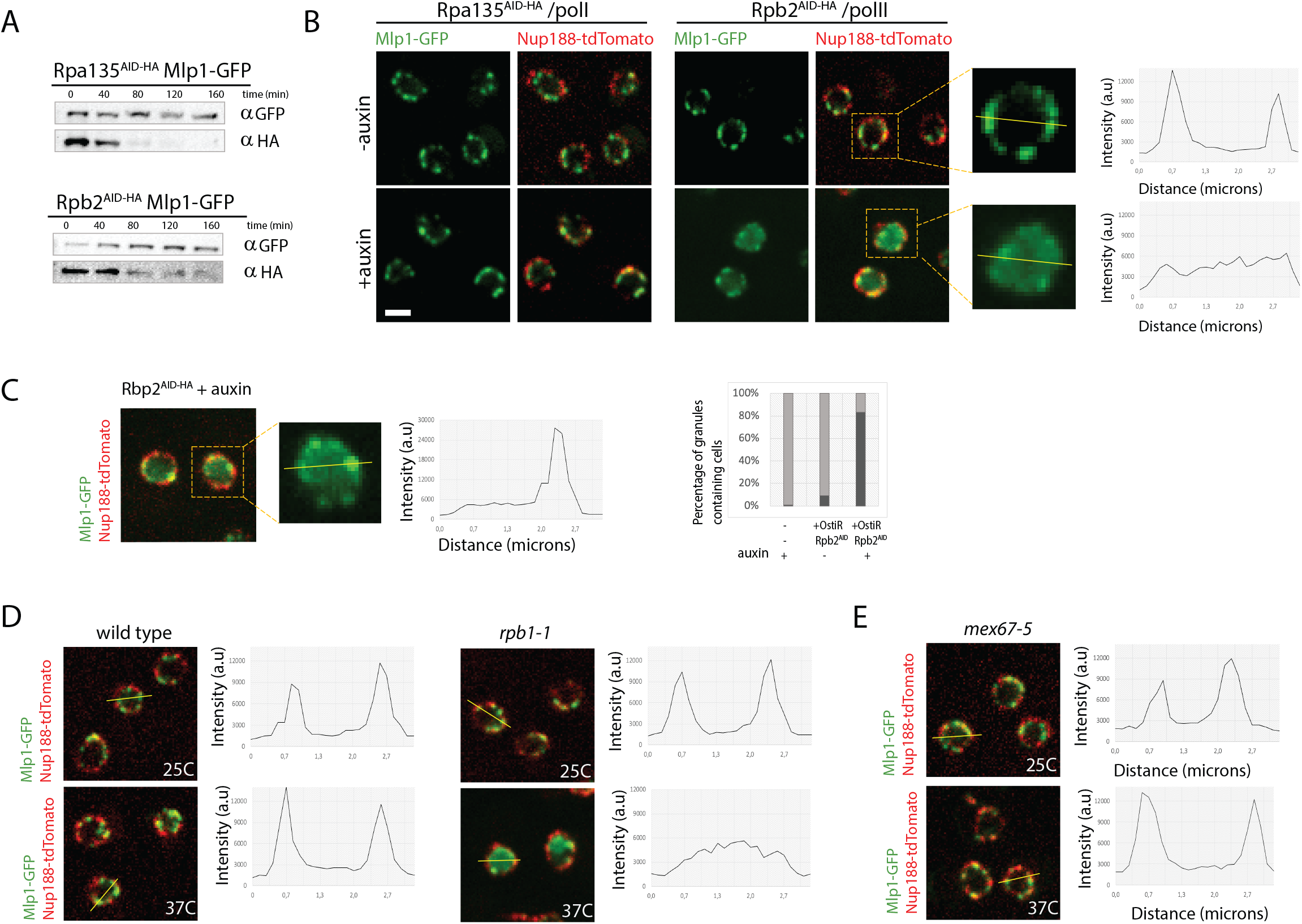
Inhibition of RNA polymerase II transcription, but not mRNA export, affects basket assembly. **A.** Western blot of total cell lysates from Rpb2^AID-HA^ and Rpa135^AID-HA^ strains at different time points upon addition of auxin. Rpb2^AID-HA^ and Rpa135^AID-HA^, and Mlp1-GFP were detected using anti-HA and anti-GFP antibodies, respectively. **B.** Fluorescent microscopy images showing Mlp1-GFP (green) localization before and 120 min after addition of 500 μm auxin in Rpb2^AID-HA^ and Rpa135^AID-HA^ strains tagged with Nup188-tdTomato (red). For the Rpb2^AID-HA^ strain, magnification of a single cell and a line scan intensity plot measuring Mlp1- GFP signal is show on the right. **C.** Image of Mlp1-GFP signal redistribution in an Rpb2^AID-HA^ strain forming a single bright Mlp1-GFP granule after 120 min of auxin, magnification of a single cell and a line plot measuring the signal intensity of Mlp1-GFP. Nup188-tdTomato signal is shown in red. Graph on the right show a quantification of cells with Mlp1-GFP granules after Rbp2 depletion. **D, E**. Mlp1-GFP distribution in wild type a *rpb1-1* and *mex67-5* tagged with Nup188-tdTomato (red) at 25°C and after shifting to 37°C. Line scan intensity plot quantifying Mlp1-GFP signal intensities are shown on the right. (Scale bar =2μm).

To help discriminate between these possibilities, we next blocked mRNA export using a temperature-sensitive mutant of the main mRNA export receptor Mex67, *mex67-5* (Santos-Rosa et al., 1998). Upon shift to non-permissive temperature (37°C), we did not observe redistribution of Mlp1 from the nuclear periphery (Figure 2E) suggesting that blocking RNA Pol II transcript production, and possibly downstream events, but not mRNA export affects basket formation. Alternatively, mRNPs reaching the periphery could be required for basket assembly.

### Rerouting of the RNA metabolism machinery induces basket formation at nucleolar pores

Similar to the compartmentalization of RNA polymerase I and II, mRNA and the components of the mRNA maturation machinery are generally restricted to the nucleoplasm and absent from the nucleolus. However, mutations in some RNA maturation and surveillance factors such as Dis3, Csl4, and the ribosome biogenesis factor Enp1 have been shown to accumulate different types of polyadenylated RNAs (rRNAs, snoRNAs, and mRNAs) and mRNP components in the nucleolus (Aguilar et al., 2020; Biplab et al., 2016). We therefore asked how a redistribution of the mRNA maturation machinery to the nucleolus affects basket formation at pores. Enp1 was depleted using a Enp1^AID-HA^ strain and basket assembly was monitored using Mlp1-GFP (Figure S4B). 120 min after addition of auxin, Mlp1-GFP signal was mostly lost from nucleoplasmic pores and redistributed to the nucleolar periphery. A similar redistribution of Mlp1-GFP was observed in Csl4 ^AID-HA^ cells upon addition of auxin (Figure S3A). This suggests that pores adjacent to nucleoli are able to assemble baskets upon redistribution of mRNPs to the nucleolus (Figure 3A) and a requirement of mRNA and/or its maturation machinery in modulating basket assembly.

**Figure 3.**
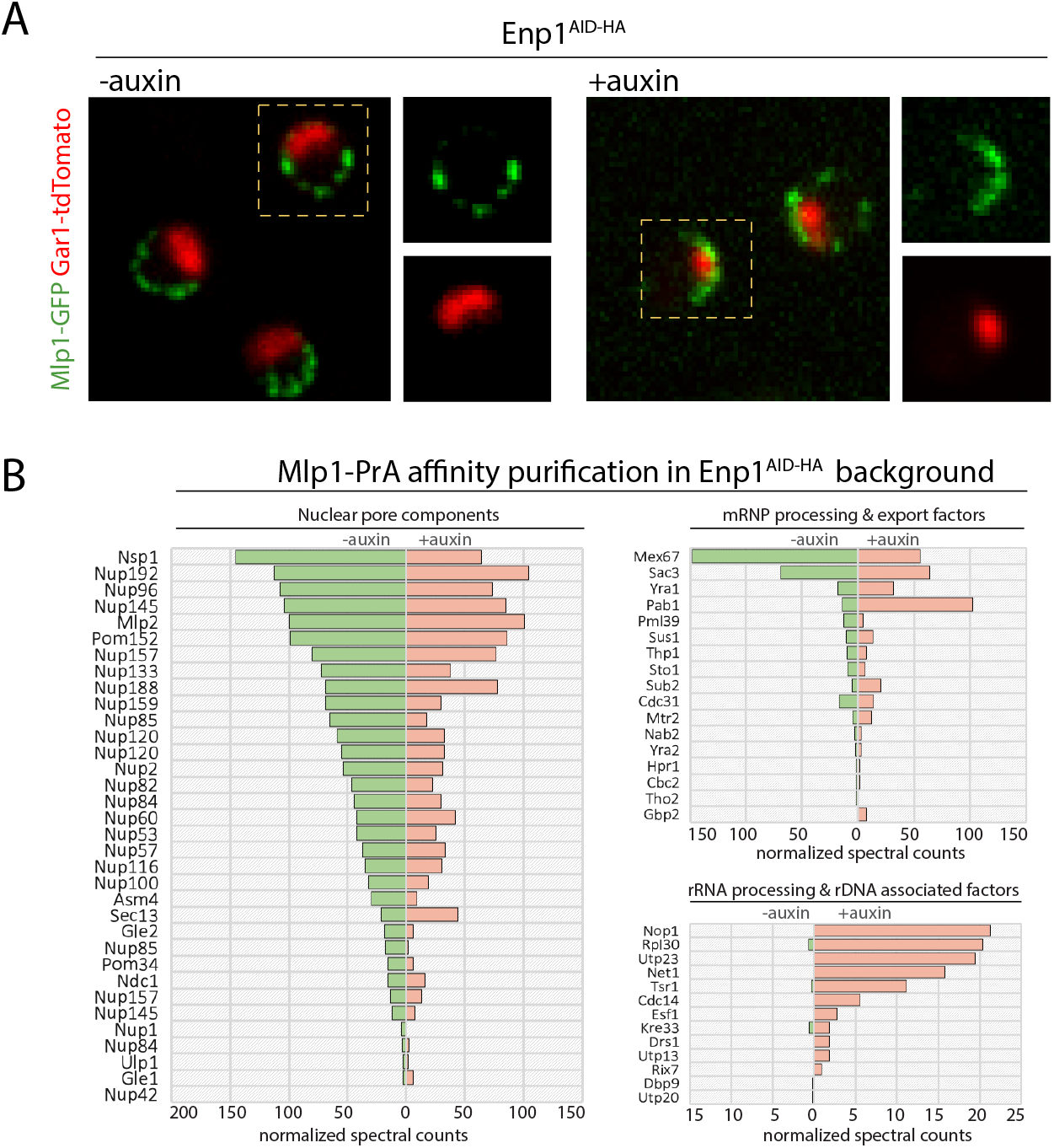
Nucleolar basket assembly upon depletion of the ribosome biogenesis factor Enp1. **A.** Mlp1-GFP (green) signal is redistributed to the nucleolar periphery in Enp1^AID-HA^ cells upon auxin treatment (120min). Nucleolus labeled by Gar1-tdTomato (red). **B.** Normalized MS spectral counts of proteins co-purified with Mlp1-PrA from Enp1^AID-HA^ cells before and after auxin treatment.

To determine that Mlp1 assembles into *bona fide* baskets on NPCs along the nucleolar periphery upon Enp1 depletion, we performed affinity purification (AP) of Mlp1-PrA and analyzed its interactome by semi-quantitative mass spectrometry (MS) with and without the addition of auxin. Under both conditions, Mlp1-associated interactomes were similar in terms of co-isolated NPC components (Figure 3B). Furthermore, consistent with accumulation of mRNPs in the nucleolus, we identified mRNA maturation and export factors associated with nucleolar Mlp1-PrA upon Enp1 depletion, in addition to low levels of nucleolar proteins. While the overall levels of mRNA maturation factors were comparable under both conditions, the poly(A)-binding protein Pab1 was significantly enriched in Mlp1-PrA APs upon addition of auxin. Moreover, under these conditions, nucleoli were often spherical, fragmented, and internalized with little or no overlap between Mlp1-GFP and Gar1-tdTomato signals (Figure S3B), reminiscent of the nucleolar invaginations caused by ectopic baskets (Figure 1C). Taken together, these data suggest that basket assembly requires elements or events of the mRNA maturation pathway that under normal conditions are restricted to the nucleoplasm, but which upon their rerouting to the nucleolus can lead to basket assembly along the nucleolar periphery. However, the assembly of nucleolar baskets appears to be incompatible with the cooccurrence of a nucleolus.

### Basket assembly requires 3’end processed and polyadenylated mRNA

To further dissect the processes along the mRNA maturation pathway required for basket assembly, we carried out a targeted AID screen, depleting selected mRNA maturation factors and monitoring Mlp1-GFP distribution upon auxin treatment. This included factors involved in co-transcriptional mRNP assembly such as components of the THO/TREX/TREX-2 complex (Yra1, Tho2, Sus1, Sac3); 5’ cap-binding (Cbc2); splicing (Prp18, Prp5, Snu17, Luc7); proteins involved in 3’ end cleavage, polyadenylation, and poly(A)-binding (Rna14, Rna15, Pap1, Nab2, Pab1); as well as proteins linked to nuclear retention of intron-containing mRNAs and quality control (Hrb1, Npl3, Gbp2, Pml1, Pml39). Depletion phenotypes were compared to depletion of Nup60, previously shown to be required for Mlp1 binding to NPCs, as well as Rpb2 and Enp1 (Figure 2, Table S1, and Figure S4B) (Lewis et al., 2007; Mészáros et al., 2015).

Phenotypes varied considerably among strains, however, shared phenotypes were observed among proteins implicated in specific stages of the pathway. Overall, depletion of early mRNP assembly factors did not affect basket formation, as neither depletion of Cbc2 nor of components of the THO/TREX complex (Yra1, Tho2) changed the Mlp1-GFP localization pattern at different timepoints after addition of auxin (Figure 4 and Table S1). Interfering with different stages of splicing did also not induce an Mlp1 relocalization as neither depletion of U1 (Luc7), U2 (Snu17) nor U5 (Prp18) snRNP-associated factors changed Mlp1-GFP localization at the nuclear periphery, with the exception of Prp5, an ATPase required for pre-spliceosome assembly (Figure 4A) (Jurica et al., 2003). Prp5 depletion resulted in an increase in nucleoplasmic Mlp1-GFP signal, similar but less pronounced to that observed in Rbp2^AID-HA^ cells after auxin treatment (Figure 2), as well as redistribution of Mlp1 along the entire nuclear periphery. In addition to its role in splicing, *prp5* mutants were shown to affect RNA Pol II transcription suggesting that this phenotype might be linked to transcription rather than splicing (Shao et al., 2020)

**Figure 4.**
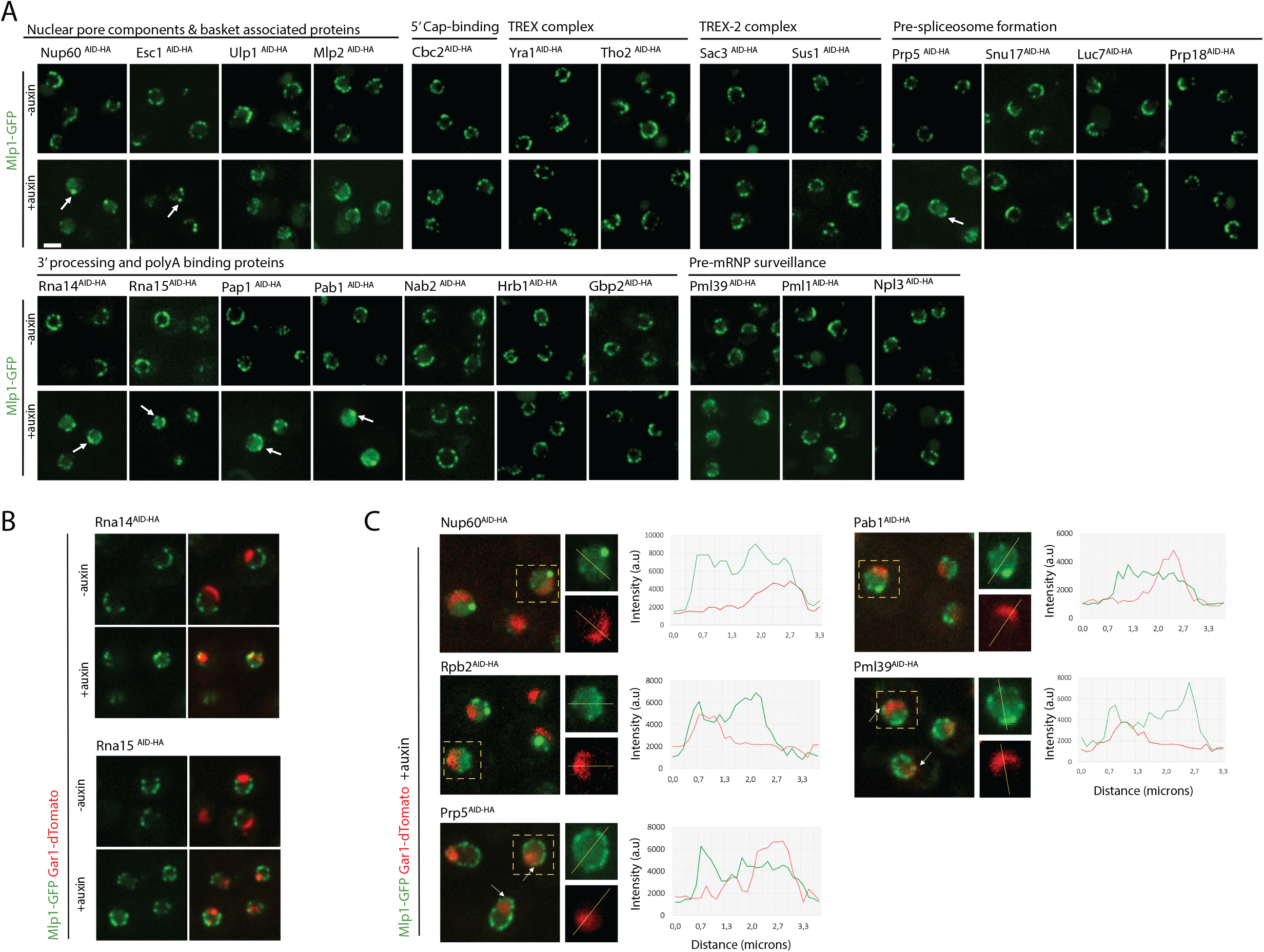
Depletion of mRNP maturation factors affecting basket assembly and Mlp1 localization. **A.** Auxin depletion screen of components along the mRNA maturation pathway and NPC associated factors. Mlp1-GFP (green) localization was scored 120min after auxin treatment. Changes in Mlp1-GFP distribution are indicated by white arrows. B. Rna14^AID-HA^ and Rna15^AID-HA^ cells show partial redistribution of Mlp1-GFP (green) to the nucleolar periphery as well as fragmented and spherical nucleoli. Nucleolus labeled with Gar1-tdTomato (red). **C.** Mlp1-GFP distribution was monitored with respect to the nucleolus in cells where baskets were destabilized upon the addition of auxin. Graphs represent the overlaps of Mlp1-GFP and Gar1-tdTomato signals (green and red curve respectively). White arrows show Mlp1-GFP signals remaining at the periphery, including in the nucleolar area after basket destabilization. (Scale bar =2μm).

Depletion of factors involved in 3’ end processing and polyadenylation, however, displayed strong Mlp1 relocalization phenotypes (Figure 4A). Specifically, depletion of factors involved in 3’ end cleavage (Rna14, Rna15), the RNA poly(A) polymerase Pap1 as well as the poly(A)-binding protein Pab1 showed altered Mlp1-GFP distribution suggesting that these steps are significant for basket formation at NPCs. Upon depletion of Rna14 and Rna15, Mlp1 was redistributed to the nucleolar periphery, similar to what we observed upon loss of Enp1 (Figure 4B and Figure 3A). In addition, these strains frequently exhibited fragmented nucleoli, a phenotype that has previously been described for *rna14-1* and *rna15-2* at non-permissive temperature (37°C) (Carneiro et al., 2007). An accumulation of poly(A) RNA in the nucleolus has also been observed in *rna14* and *rna15* mutants as a consequence of disrupting 3’end cleavage, polyadenylation, and export (Brodsky et al., 2000; Casolari et al., 2004; Dunn et al., 2005). Similar to *csl4* and *enp1* mutants, where nucleolar accumulation of poly(A) RNA results in nucleolar sequestration of mRNPs, redistribution of the mRNA maturation machinery to nucleoli might be responsible for basket formation along the nucleolar periphery in Rna14^AID-HA^ and Rna15^AID-HA^ cells (Aguilar et al., 2020). While Pap1 depletion also led to a redistribution of Mlp1, an increase in nucleoplasmic Mlp1 levels and decrease of Mlp1 signal along the nuclear periphery was observed, along with concomitant formation of larger Mlp1 foci in ~50% of the cells, reminiscent of a Rbp2 depletion phenotype (Figure 4C). A similar but somewhat stronger phenotype was seen upon loss of Pab1 (Figure 4A, C), which also resembled that of a Nup60 depletion strain. No change in Mlp1 pattern was observed upon depletion of the Mlp1 interactor and poly(A) binding protein Nab2, nor upon that of Gbp2 or Hrb1, two poly(A)-binding proteins linked to the maturation of intron-containing mRNAs, or Npl3 (Figure 4A).

We also tested proteins that have previously been linked to nuclear basket function for a role in Mlp1 localization. Of the two pre-mRNA surveillance factors (Pml1, Pml39), only Pml39 displayed a phenotype upon auxin treatment, where Mlp1 signal was decreased along the nuclear periphery with an increase in nucleoplasmic Mlp1 away from NPCs. On the other hand, TREX-2 components (Sac3, Sus1) showed no phenotype (Figure 4A). We furthermore determined Mlp1-GFP localization upon depletion of NPC- and nuclear basket-associated factors Esc1, Ulp1, and the Mlp1 paralog Mlp2. Upon addition of auxin, Esc1^AID-HA^ cells exhibited formation of larger Mlp1 granules along the nuclear rim with a simultaneous loss of evenly distributed Mlp1 rim staining, while Ulp1 and Mlp2 depletion showed no phenotype (Figure 4A).

Overall, we did not observe a common phenotype, suggesting that basket assembly is not a consequence of nuclear mRNA metabolism per se but rather the completion of specific steps. Mlp1-GFP granules observed upon depletion of Rpb2, Pab1, Pap1, Nup60, and Esc1 were similar to those previously described in this work insofar as they were systematically excluded from nucleolar areas (Figure 1C, Figure 4C and Figure S4A). On the other hand, when nucleoplasmic Mlp1 levels were significantly increased upon auxin treatment and baskets disassembled (i.e., Rpb2^AID-HA^, Prp5^AID-HA^ Pab1^AID-HA^, Pml39^AID-HA^ Nup60^AID-HA^), nuclear Mlp1-GFP signal overlapped with the nucleolar region corroborating our previous observation that the nucleolus does not present a diffusion barrier for free-diffusing Mlp1 (Figure 1E; Figure 4C). These observations suggest that Mlp1 exclusion from the nucleolar phase occurs upon Mlp1 multimerization during nuclear basket formation or granule assembly. Moreover, in Prp5, Pml39, and to a lesser extend in Pab1, Pap1, and Rpb2 depleted cells, we observed a weak Mlp1-GFP signal at the nuclear periphery, as well as along the nucleolus:nuclear envelope interface upon Prp5, Pab1, and Pml39 depletion (Figure 4C, white arrows). This Mlp1-GFP signal along the nucleolar periphery was not associated with any nucleolar fragmentation or invagination as was observed upon loss of Enp1 or Csl4 (Figure S3B), or ectopic basket formation (Figure 1C). This may indicate that depletion of Pap1, Pab1, Pml39 or Prp5 does not affect the ability of Mlp1 to bind an NPC per se, unlike in Nup60-depleted cells (Mészáros et al., 2015), but rather their capacity to form fully assembled baskets.

Taken together, our results suggest that basket formation at NPCs is a consequence of completing specific steps of nuclear mRNA maturation and requires the presence of poly(A) transcripts as we observed a significant loss of Mlp1-GFP along the nuclear periphery and its redistribution into the nucleoplasm upon depletion of Rbp2, Prp5, Pap1, Pab1, and Pml39. Yet not all poly(A)-binding proteins appeared to be required for basket formation as neither depletion of Gbp2, Hrb1 nor Nab2, which has been linked to Mlp1 in the context of mRNA export and shown to directly interact with the Mlp1 C-terminal region (Green et al., 2003), affected nuclear basket formation.

### Not all nucleoplasmic pores contain baskets

The requirement for an active mRNP maturation pathway for basket formation might suggest that baskets assemble randomly at individual NPCs, and that not all pores assemble baskets at all times or with the same frequency, which could result in a functional heterogeneity at nucleoplasmic NPCs. Consistent with such a model, Mlp1-GFP distribution at the nuclear periphery, imaged using spinning disk confocal microscopy, often showed a discontinuous staining pattern (Figure 1A). To investigate possible NPC heterogeneity at nucleoplasmic pores, we analyzed distribution and co-localization of different NPC components using Structured Illumination microscopy (SIM) (Figure 5). Signal distribution of two components of the central framework Y-complex, Nup84-GFP and Nup188-tdTomato, serving as controls, showed an overall continuous staining and co-localization pattern along the nuclear periphery. Quantification of normalized Nup84-GFP and Nup188-tdTomato signal intensities did not reach background intensity levels, consistent with a homogeneous distribution of NPCs all around the nuclear periphery (Figure 5A) (Steinberg et al., 2012; Winey et al., 1997). Mlp1-GFP staining, however, showed a discontinuous distribution with regions where GFP signal was interspaced by segments devoid of Mlp1-GFP staining, suggesting that several regions of basket-less pores occupy the nuclear periphery.

**Figure 5.**
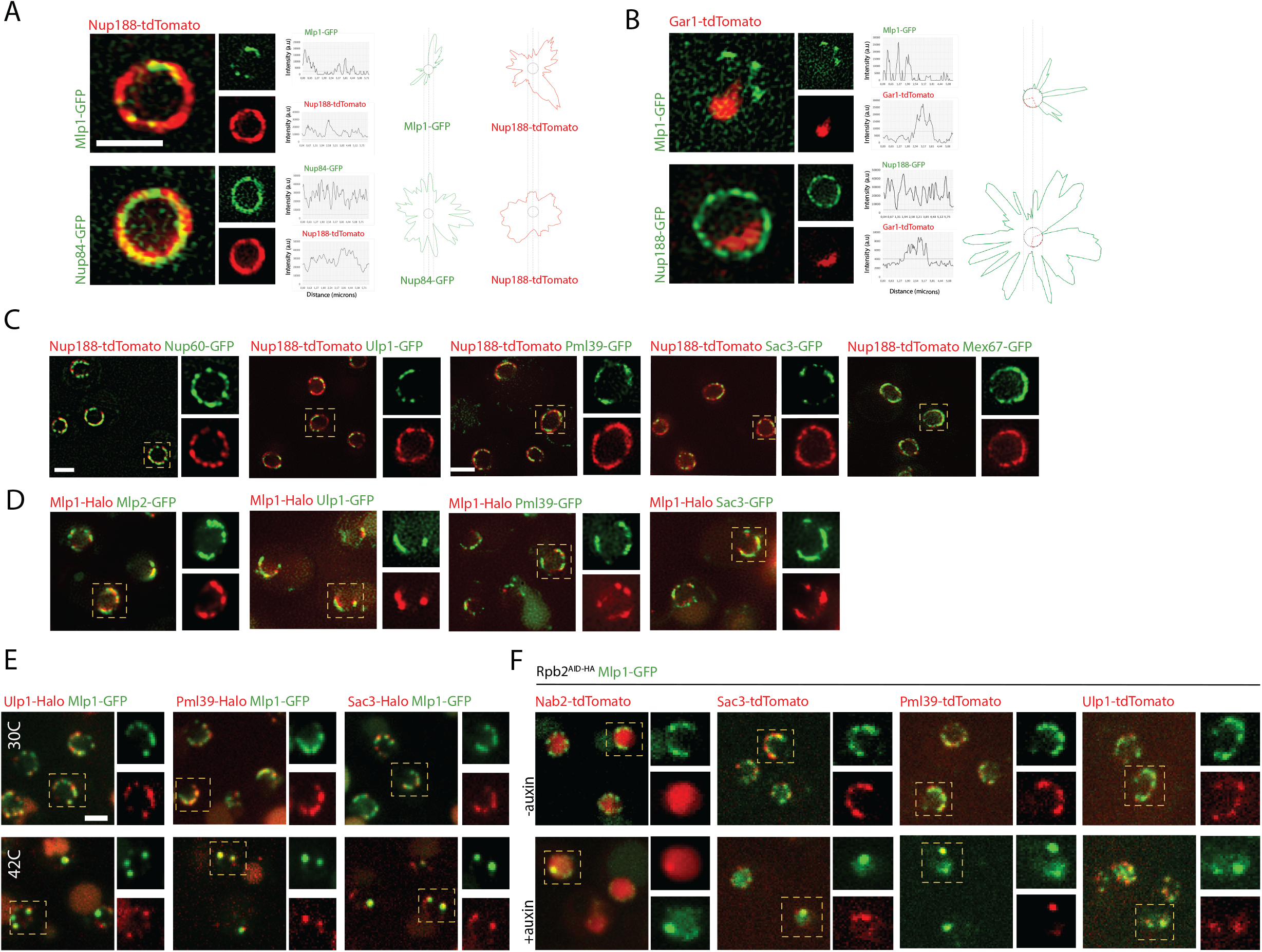
Baskets assemble on a subset of NPCs in the nucleoplasm and co-localize with select proteins at the nuclear periphery. **A.** Structured illumination microscopy (SIM) images of cells double tagged for Nup188-tdTomato (red) and either Mlp1-GFP or Nup84-GFP (green) and respective signal intensities along the nuclear periphery are shown on the right. Grey circles represent average background values for each signal. **B.** SIM images of cells double tagged for nucleolar Gar1-tdTomato (red) and either Mlp1-GFP or Nup188-GFP (green) and respective signal intensities along the nuclear periphery are shown on the right. Grey circles represent average background values for each signal. **C.** SIM co-localization analysis of cells double tagged for Nup188-tdTomato (red) and select GFP-tagged proteins (green). **D.** SIM colocalization analysis of cells double tagged for Mlp1-Halo (red) and select GFP-tagged proteins (green). **E.** SIM co-localization analysis of cells double tagged for Mlp1-GFP (green) and select Halo-tagged proteins (red) at 30°C or 42°C. **F.** SIM co-localization analysis of cells double tagged for Mlp1-GFP (green) and select Halo-tagged proteins (red) in Rbp2^AID-HA^ cell with and without auxin depletion. (Scale bar =2μm)

To better define the position of basket-less nucleoplasmic regions in relation to the nucleolus, we imaged double-tagged Mlp1-GFP / Gar1-tdTomato cells. In addition to the nucleolar area, several regions along the nucleoplasmic periphery exhibited a lack of Mlp1-GFP, indicating that not all nucleoplasmic pores contain baskets (Figure 5B). Previous works have shown that basket proteins Nup60, Nup1, and Nup2 are present on pores adjacent to nucleoli, and co-staining of Nup-188-tdTomato with Nup60-GFP confirmed association of Nup60 with all NPCs (Figure 5C and Figure S5A) (Galy et al., 2004). However, Mlp2-GFP showed a localization pattern similar to that of Mlp1 and co-localized with Mlp1-Halo (Figure 5D), in line with the requirement of Mlp1 for Mlp2 perinuclear localization (Palancade et al., 2005). Moreover, this indicates that NPC heterogeneity implicates Mlp1/2, but not the other asymmetric nuclear basket proteins, Nup60, Nup1, or Nup2. Together, our data show that NPC heterogeneity extends beyond the nucleolus and implies that basket formation is not a default state of nucleoplasmic NPCs.

### Specific nuclear mRNA maturation factors are enriched at basket-containing NPCs

Various proteins generally not considered *bona fide* nucleoporins associated with NPCs have been linked to the nuclear basket, including components of the TREX-2 complex, Pml39, and the ubiquitin-like modifier Ulp1, and were previously shown to be excluded from the nucleolar periphery (Bonnet et al., 2015; Zhao et al., 2004). Thus, we analyzed the distribution of the TREX-2 main scaffold protein Sac3-GFP, Ulp1-GFP and Pml39-GFP relative to Nup188-tdTomato. All three proteins displayed an uneven pattern along the periphery similar to Mlp1-GFP staining (Figure 5C). Moreover, Ulp1-GFP, Sac3-GFP and Pml39-GFP co-localized with Mlp1-Halo signal in double-tagged cells, suggesting that these proteins associate preferentially with baskets-containing nucleoplasmic NPCs (Figure 5D and Figure 5B). Moreover, these proteins also colocalized in Mlp1 granules that formed upon heat-shock at 42°C (Figure 5E), and Sac3 was co-isolated with Mlp1 at nucleolar NPCs formed upon Enp1 depletion (Figure 3B), further linking these proteins to basket-containing NPCs.

The localization of these proteins, however, is different to that of Mex67-GFP, which showed a distribution pattern that occupied the entire nuclear periphery, overlapping with Nup188-tdTomato (Figure 5C). This is consisted with recent models that suggest that Mex67 is a *bona fide* nuclear pore component associating with pores independent of its association with RNPs (Derrer et al., 2019). Moreover, it may also indicate that all NPCs are, in principle, able to export mRNAs using the Mex67-dependent RNA export pathway, but that some are further functionalized by the presence of a basket and associated factors implicated in mRNA metabolism (e.g., TREX-2, Ulp1 and Pml39).

To further determine whether the presence of baskets on nuclear pores correlates with the presence of a specific basket protein interactome, we characterized the interactomes of NPCs with and without nuclear baskets by AP-MS. First, to analyze the general nuclear pore interactome (‘all pores’), we carried out single-step affinity purifications (ssAP) of nuclear pores followed by mass spectrometry from an Mlp1-PrA/Nup133-GFP double-tagged yeast strain, using Nup133-GFP as bait protein (Figure S6A) (Trahan et al., 2016). To ensure the capture of dynamic interactors such as Mlp1, we stabilized NPCs and associated proteins using a short in-lysate glutaraldehyde fixation prior to incubation with antibody-conjugated magnetic resin (Subbotin et al., 2014). To compare the interactomes of pores with and without nuclear baskets, we applied a differential affinity purification approach (dAP) that enabled us to separate and isolate the two types of pores from the same lysate via two consecutive APs. In a first step, incubation with IgG-conjugated resin allows for the isolation of Mlp1-PrA and its associated complexes, including basket-containing nuclear pores (‘Basket^plus^’) (Figure S6A); a second affinity purification was carried out on the flow-through using Nup133-GFP to isolate the remaining basket-less pores (‘Basket^minus^’). Moreover, to identify proteins associated with nuclear pores in an Mlp1- and/or poly(A) RNA-dependent manner, we also affinity purified NPCs and their interactome from a *Δmlp1/2*/Nup133-GFP cells and Enp1^AID-HA^ cells upon auxin treatment using Nup133-GFP and Mlp1-PrA, respectively. AP-MS and dAP-MS experiments were carried out in triplicate and normalized across samples (see Tables S5 and S6). While for our analysis Mlp1-PrA complexes were considered NPC-associated, we cannot rule out that some proteins identified with Mlp1-PrA interact only transiently with the periphery or with free nucleoplasmic Mlp1.

AP-MS data for ‘all pores’ (Nup133-GFP) revealed an enrichment for nucleoporins, which constituted the majority of the isolated NPC interactome (77%), proteins involved in mRNA export and processing (7%), proteasome components (5%), ribosome biogenesis factors (5%), karyopherins (3%), transcription and chromatin-associated proteins (2%) as well as spindle pole body (SPB) proteins, spliceosome components, surveillance factors and other nucleolar proteins (~1%)(Figure 6A). Efficient separation of basket-containing (‘Basket^plus^’, Mlp1-PrA) and basket-less pores (‘Basket^minus^’, Nup133-GFP) was confirmed by a significant decrease of Mlp1 and Mlp2 levels in ‘Basket^minus^’ samples compared to ‘all pores’ (Figure 6B).

**Figure 6.**
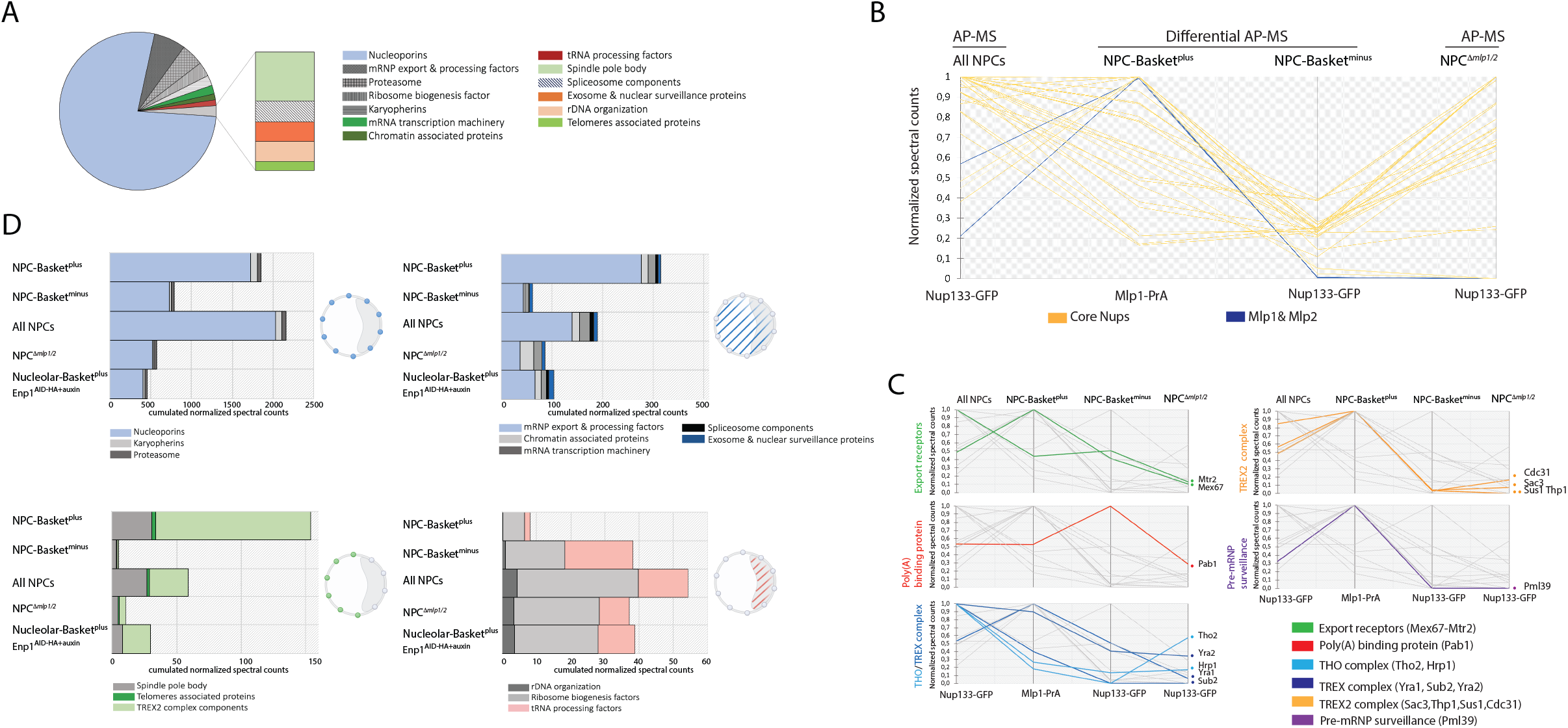
Dissection of differential NPC interactomes. **A.** Overview NPC interactome from the total pore AP. Each protein categories represent the sum of the normalized spectral counts of proteins purified with NPCs. **B.** Normalized spectral counts values for nuclear pore and basket proteins affinity purified from either ‘All NPC’, Mlp1-PrA (Basket^plus^), and Nup133-GFP (Basket^minus^) using Mlp1-PrA/Nup133-GFP or *Δ mlp1/2/*Nup133- GFP cells are presented on a scale from 0 to 1. Each line represents the relative abundance for one protein between different APs. **C.** Histograms representing the sum of normalized spectral counts of proteins co-purified with either ‘All NPC’, Mlp1-PrA (Basket^plus^), and Nup133-GFP (Basket^minus^) from Mlp1-PrA/Nup133-GFP, *Δ mlp1/2/* Nup133-GFP cells and Mlp1- PrA/Nup133-GFP/Enp1^AID-HA^ cells upon auxin treatment, and analyzed according to function in RNA metabolism across different subnuclear compartments: at the nuclear periphery (top left); in the nucleus (top right); at the nuclear periphery but excluded from the nucleolus (bottom left); in the nucleolus (bottom right). Cartoons represent yeast nuclei with the nucleolus outlined as grey crescent. Spheres represent proteins associating with the nuclear periphery without bias in blue and outside the nucleoli in green. Dashed blue lines represent nuclear proteins with no reported nucleolus/nucleoplasm bias. Dashed red lines represent nucleolar proteins. **D.** Normalized spectral counts values for proteins involved in nuclear mRNA metabolism affinity purified from either ‘All NPC’, Mlp1-PrA (Basket^plus^), and Nup133-GFP (Basket^minus^) using Mlp1- PrA/Nup133-GFP or *Δ mlp1/2/*Nup133-GFP cells are presented on a scale from 0 to 1. Each line represents the relative abundance for one protein between different APs. See also Tables S5 and S6.

We then analyzed the relative abundance of RBPs as well as factors required for mRNA maturation, mRNA export and surveillance by comparing their normalized spectral counts across the different samples (Figure 6C, D; Figure S6B, C). The mRNA export receptor heterodimer Mex67:Mtr2 was identified with both basket-containing (‘Basket^plus^’) and basket-less (‘Basket^minus^’) pores as well as with NPCs in *Δmlp1/2* cells, consistent with Mex67 localization (Figure 5C), supporting the notion that Mex67 is systematically present at NPCs (Derrer et al., 2019) and, moreover, that RNP export may occur through both types of pores. Yra1, other THO/TREX complex components as well as the poly(A)-binding protein Pab1 were also purified with both basket-containing (‘Basket^plus^’) and basket-less (‘Basket^minus^’) pores and with NPCs in *Δmlp1/2* cells (Figure 6C; Figure S6B) suggesting that mRNPs containing these factors can interact with NPCs independent of a basket or Mlp1 and is consistent with previous observations that deletion of MLP1/2 causes only a moderate mRNA export defect (Galy et al., 2004; Green et al., 2003). These proteins, however, were all enriched with Mlp1-PrA in Enp1^AID-HA^ cells upon auxin treatment (Figure S6B) indicating that while, unlike Pab1, THO/TREX components are not required for basket formation (Figure 4A), they are tightly linked to nuclear poly(A) mRNA metabolism (Aguilar et al., 2020). A similar behavior was also observed for the poly(A)-binding proteins Nab2 and Gbp2, yet their overall spectral counts were too low to compare their distribution across samples (Figure S6B).

Unlike THO/TREX, TREX-2 components (Sac3, Sus1, Thp1, Cdc31) were enriched with basket-containing (‘Basket^plus^’) over ‘all pores’ and absent in basket-less (‘Basket^minus^’) pores, while present at low levels in *Δ mlp1/2* cells (Figure 6C; Figure S6B). These results are consistent with our previous observation that Sac3-GFP co-localized with Mlp1-Halo along the nuclear periphery (Figure 5C) and the suggestion that TREX-2 associates preferentially with baskets-containing nucleoplasmic NPCs. While Ulp1 levels were too low to compare its distribution across samples, similar to TREX-2, the pre-mRNA surveillance factor Pml39 was significantly enriched with basket-containing (‘Basket^plus^’) pores (Figure 6C; Figure S6B), again in agreement with our observation that Pml39 localizes only with Mlp1-Halo and basket-containing pores (Figure 5C). Pml39 was significantly decreased in *Δ mlp1/2* cells, while it was enriched with Mlp1-PrA at nucleolar pores in Enp1^AID-HA^ cells, both consistent with previous data that its association with the NPC is dependent on the presence of a nuclear baskets, i.e., Mlp1 (Palancade et al., 2005).

To further analyze the differences between basket-containing and basket-less pore interactomes, we organized identified proteins based on their sub-localization within the nucleus. To that end, co-purified proteins were divided into four groups (Figure 6D): (i) nucleoporins and NPC-associated proteins present in both nucleus and nucleolus (upper left); (ii) nuclear non-periphery-associated proteins involved in RNA metabolism (upper right); (iii) complexes associated with nucleoplasmic pores (lower left); and (iv) nucleolar proteins (lower right). Proteins from groups i and ii were identified across all samples, consistent with their relative abundance compared to ‘all pores’ (Figure S6B). Again, while many mRNA maturation and export factors (group ii) were found enriched with Mlp1-PrA and ‘Basket^plus^’ pores, their presence with ‘Basket^minus^’ pores and NPCs in *Δ mlp1/2* cells indicates that these interactions are not basket-dependent. Yet, the significantly lower number of mRNA maturation and export factors found with NPCs in the *Δ mlp1/2* background the basket may facilitate these interactions.

Within group iii, complexes known to interact with nucleoplasmic pores, possibly in a basket-dependent manner (e.g., TREX-2, SPB proteins), were significantly enriched with basket-containing (‘Basket^plus^’) in wild-type and Enp1^AID-HA^ cells and mostly absent from basket-less pores (‘Basket^minus^’). Conversely, group iv nucleolar proteins were found significantly underrepresented with nucleoplasmic basket-containing pores (‘Basket^plus^’) compared to basket-less pores (‘Basket^minus^’) and ‘all pores’ (Figure 6D). This category was also slightly increased with nucleolar basket-containing NPCs upon relocalization of Mlp1 to the nucleolar periphery in Enp1^-^depleted cells suggesting overall that while pre-ribosomes and tRNPs may preferentially interact with basket-less (‘Basket^minus^’) pores, but may associate with basket-containing pores (‘Basket^plus^’) under specific conditions. However, as no ribosome export factors were identified in our AP-MS samples, and depletion of Enp1 is a terminal phenotype, it is unlikely that these pores are actively exporting pre-ribosomes or mRNPs (Aguilar et al., 2020); alternatively, these factors, except for Mex67, may be more transiently associated with NPCs.

Taken together, our AP-MS results show that, overall, basket-less and basket-containing pores associate with a shared set of mRNA maturation and export factors, suggesting that mRNPs can bind, and most likely be exported, through both types of pores. This is also in accordance with the observation that mRNA export does not require Mlp1/2, or a nuclear basket (Powrie et al., 2011). However, despite that, a number of specific factors, namely TREX-2 components and the pre-mRNA surveillance factor Pml39, were significantly enriched with Mlp1-PrA and basket-containing pores suggesting that these pores may represent a differentially regulated export route for at least a subset of mRNPs. Finally, these AP-MS data link basket formation to overall poly(A)-RNA metabolism, as a large number of mRNA maturation factors associated with nucleolar Mlp1 and basket-containing NPCs upon sequestration of poly(A) transcripts to the nucleolus.

### A distinct Mlp1 mRNA interactome suggests differential export via basket and basket-less pores

Interactome dissection of NPCs with and without basket (i.e., Mlp1) showed that while mRNP-associated factors are found with all pores, there was a distinct subset of mRNP export and processing factors enriched with Mlp1-PrA, suggesting that some mRNAs may be preferentially exported via basket-containing pores. To determine if subsets of nuclear mRNAs are differentially associated with nucleoplasmic basket-containing pores (‘Basket^plus^’) or basket-less pores (‘Basket^minus^’), AP-RNA-seq was performed on oligo-dT-purified RNA samples isolated with either ‘all pores’ (Nup133-GFP), or basket-containing pores (‘Basket^plus^’) and basket-less pores (‘Basket^minus^’) using differential APs as described above. Control experiments were carried out using strains expressing Protein A or GFP alone, and RNAs identified in these samples were considered background and filtered from those identified with Mlp1-PrA and Nup133-GFP, respectively. Following background removal, the remaining RNAs in each sample were then compared against a poly(A)-RNA library generated from a total RNA (i.e., the cellular transcriptome) to identify transcripts differentially associated with pores, using a log2 fold change (FC) cut-off of greater than (>) 1.

Using this approach, we found 680 transcripts enriched with ‘all pores’ (Nup133-GFP), 1379 with ‘Basket^plus^’(Mlp1-PrA), and 484 with ‘Basket^minus^’ (Nup133-GFP) pores (Figure 7A, left). Comparing these, 746 transcripts were found exclusively enriched with ‘Basket^plus^’, while only five were exclusively enriched with ‘Basket^minus^’ pores (Fig7.A, left), which may indicate a preferential association of a subset of transcripts with basket-containing pores. To explore this idea, the log_2_FC values for the 439 mRNAs enriched in both the ‘Basket^plus^’ and ‘Basket^minus^’ samples were compared, and it was found that log2FC values were consistently higher in ‘Basket^plus^’ samples (paired t-test, estimate = 1.22, p < 0.001) (Figure 7A, right). In other words, although these 439 mRNAs can be identified in both samples, their enrichment is greater in basket-containing pores. These data suggest a preferential association of transcripts with basket-containing pores, or possibly, the association of Mlp1 with transcripts in the nucleoplasm prior to their association with the NPC.

**Figure 7.**
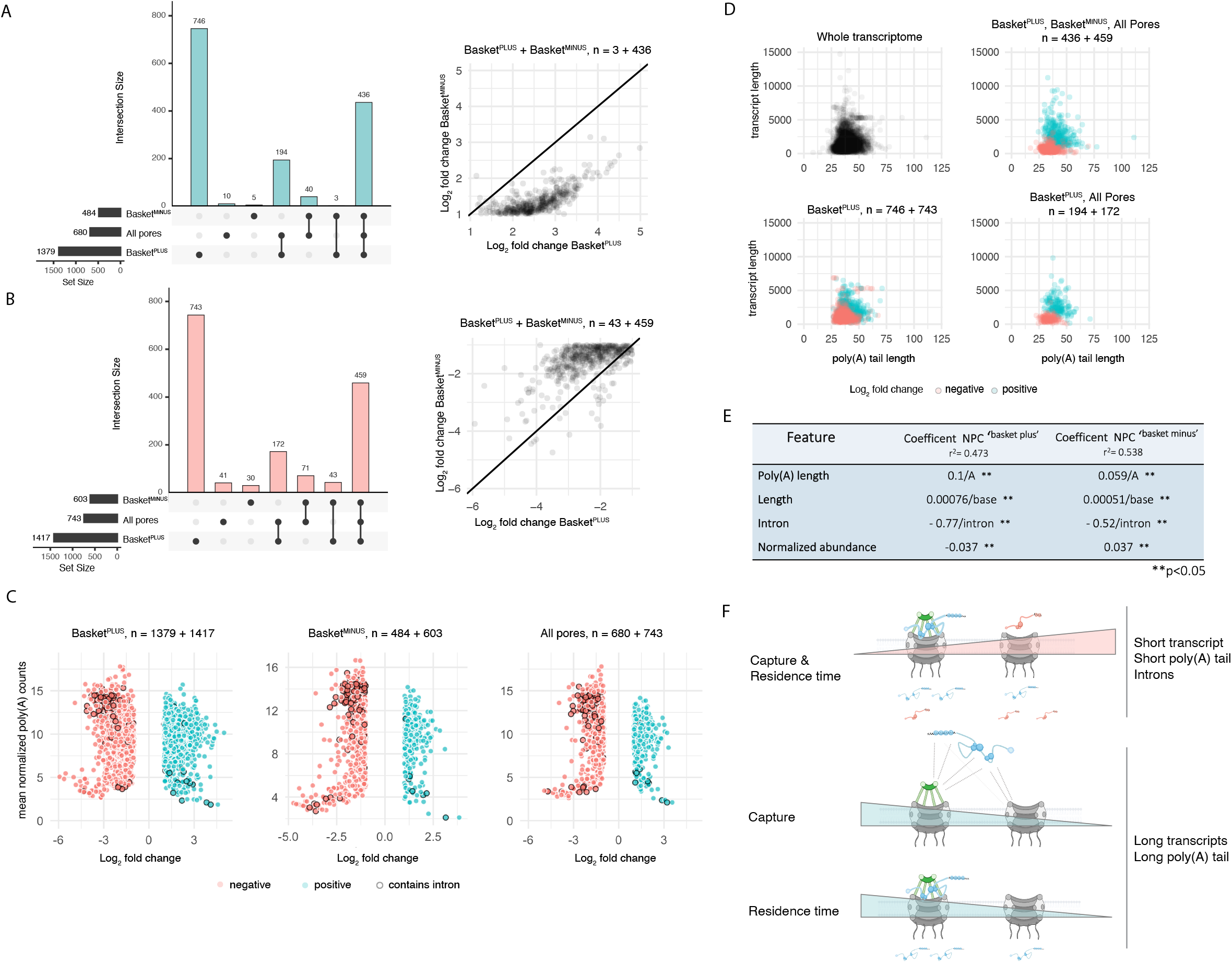
A differential Mlp1 RNA interactome suggests selective transport. **A.** Upset plot. Bar graph on the bottom left represents transcripts identified with ‘All NPCs’, and differential ‘Basket^plus^’ and ‘Basket^minus^’ NPC APs with a log_2_FC > 1 over poly(A) libraries. Bar graph on top represents the transcripts exclusively enriched. Bar graphs represent intersection sizes while dots in line with bars indicate shared transcript sets. Scatter plot on the right shows transcripts that are enriched in both ‘Basket^plus^’ and ‘Basket^minus^’ samples and have higher log_2_FC values in ‘Basket^plus^’ compared to ‘Basket^minus^’. **B.** Upset plot on the left represents the transcripts significantly underrepresented with a log_2_FC < 1 over poly(A) libraries across samples. Bar graph on top represents the transcripts exclusively underrepresented. Bar graphs represent intersection sizes while dots in line with bars indicate shared transcript sets. Scatter plot on the right shows transcripts that are depleted in both ‘Basket^plus^’ and ‘Basket^minus^’ have lower log_2_FC values in ‘Basket^plus^’ compared to ‘Basket^minus^’. **C.** Scatter plots of log_2_FC values against mean normalized transcript expression in the poly(A) library. Plots are labelled by differential AP and intersection numbers from the upset plots. **D.** Scatter plots of poly(A) tail length against transcript length. Plots are labelled by differential AP (dAPs) and intersection numbers from the upset plots. Transcripts enriched in dAPs tend to have longer poly(A) tails and/or longer transcript length compared to transcripts that are underrepresented. **E.** Linear regression models built from poly(A) tail length, transcript length, intron presence/absence and mean poly(A) normalized transcript counts for ‘Basket^plus^’ and ‘Basket^minus^’ samples. All coefficents were significant (p < 0.05). **F.** Cartoons illustrating models of mRNP capture and residence time at ‘Basket^plus^’ and ‘Basket^minus^’ pores based on features such as transcript and poly(A) tail lengths and presence or absence of introns.

Conversely, we identified 743 transcripts underrepresented with ‘all pores’, 1417 transcripts with ‘Basket^plus^’, and 603 with ‘Basket^minus^’ (Figure 7B, left). Among these, 743 mRNAs were specifically depleted from ‘Basket^plus^’ samples and only 30 from ‘Basket^minus^’ (Figure 7B, left). When considering log2FC values, the vast majority of the 502 mRNAs underrepresented across ‘Basket^plus^’ and ‘Basket^minus^’ samples had lower log_2_FC values in basket-less pores compared to basket-containing pores (paired t-test, estimate = −0.85, p < 0.001) (Figure 7B, right). Overall, this suggests that these transcripts may form unstable interactions with basket-containing pores, have shorter residence times at the NPC, are exported more rapidly, or may indicate a preferential association of those transcripts with basket-less pores.

Gene ontology (GO) enrichment analyses did not reveal any significant bias in the functional classification of the different transcripts co-purified in ‘Basket^minus^’ and ‘Basket^plus^’ samples. Therefore, we next used normalized counts across our AP data to test whether the relative enrichment of each transcript across samples was related to cellular transcript abundance. Overall, we did not observe a strong correlation between transcript abundance in cells and their relative enrichment at pores (Figure 7C; Figure S7A). However, transcripts generated from intron-containing genes, which are among the most highly expressed transcripts, were underrepresented across all three samples (Figure 7C; Figure S7A), suggesting that mRNA interaction with pores may vary depending on specific features. To identify other features that may influence the relative enrichment of transcripts with pores, we further considered transcript length and poly(A) tail length, using published poly(A) sequencing data (Tudek et al., 2021). We observed that transcripts enriched across all samples tended to be longer and to have longer poly(A) tails compared to underrepresented transcripts (Figure7D; Figure S7B, C), suggesting that mRNA length as well as the length of the poly(A) tail may influence the interaction of mRNAs with NPCs. This distinction was not observed in control samples, indicating that this relationship is not the result of an experimental bias related to mRNA purification or sequencing (Figure S7B, C).

To further examine the impact of transcript length, poly(A) tail length, presence of an intron, and transcript abundance on the association of mRNAs with ‘Basket^plus^’ and ‘Basket^minus^’ pores, we utilized linear regression to model the relationship between these variables and log2FC (enrichment) in the dAPs. The resulting models show that presence of introns negatively influenced the enrichment of transcripts with NPCs (Figure 7E) and the penalty for the presence of an intron was greater for mRNAs associating with ‘Basket^plus^’ than ‘Basket^minus^’ pores. The model further confirmed that transcript length and poly(A) tail length had a positive influence on transcript enrichment for both ‘Basket^minus^’ and ‘Basket^plus^’ pores, which was larger for mRNAs found associated with basket-containing pores (Figure 7E). This data together suggests that longer transcript and poly(A) tail lengths favor mRNA::NPC association at different types of pores, while the presence of an intron makes it less likely to observe an interaction. Interestingly, the log2fc increase per base was higher for the poly(A) tail length than for the total transcript length, indicating increased poly(A) tail length to be more significant than overall transcript length.

Taken together, our results suggest a preferential association of mRNAs with basket-containing and basket-less pores based on features such as poly(A)tail length, transcript length and the presence of introns. This may be reflective of differential export dynamics for mRNAs/mRNPs. Considering that, in yeast, intron-containing transcripts are generally short (<1000nt), their short size and/or presence of an intron may lead to differential interactions with pores and faster export kinetics as suggested by their general under-representation across all NPCs (Figure 7F, i). Conversely, longer mRNAs were observed enriched across all pores, suggesting slower export kinetics or longer residence times, possibly due to necessary mRNP rearrangements prior to export (Figure 7F, ii). Notably, longer transcript and poly(A) tail lengths favored basket-containing pores/Mlp1, suggesting a potential role for the basket in directing the export of a subset of mRNPs with distinct features.

## DISCUSSION

The nuclear basket is generally considered an integral part of the NPC, thought to modulate many functions, including providing a first point of contact for exporting mRNPs at the nuclear pore, as well as a staging platform for quality control. In this study, we show that in *S. cerevisiae* the nuclear basket is not a default structure of all pores. While it has previously been observed that in yeast nuclear pores adjacent to the nucleolus are devoid of a nuclear basket, our findings demonstrate that even in the nucleoplasm baskets are not present on all NPCs. Basket assembly is linked to mRNA metabolism as inhibition of RNA polymerase II transcription results in the abrogation of basket assembly at nucleoplasmic NPCs. Specifically, interference with 3’ end processing and polyadenylation also leads to loss of nuclear baskets, linking specific steps of mRNA maturation to basket assembly. Unexpectedly, depletion of Pab1, but not Nab2 which has been closely linked to Mlp1 and the basket, affected basket assembly at the periphery. Moreover, our proteomic and microscopy data reveal that these nucleoplasmic baskets associate a specialized interactome, while AP-RNA-seq analysis suggests a preferential association of long, but not intron-containing mRNAs, with basket-containing pores. Taken together, our data points towards nuclear pore heterogeneity and an RNA-dependent mechanism for functionalization of nuclear pores in budding yeast through nuclear basket assembly.

### Basket assembly is driven by early steps in nuclear mRNA metabolism

The nuclear basket in budding yeast has long been connected to mRNA quality control and export (Strambio-De-Castillia et al., 2010). Studies investigating the nuclear retention of inefficiently spliced pre-mRNAs found increased leakage of unspliced transcripts to the cytoplasm in the absence of Mlp1and Mlp2 (Galy et al., 2004). Moreover, genetic studies have linked the basket to the RNA exosome, further underlining its connection to nuclear (pre-)mRNA surveillance (Vinciguerra et al., 2005). While Mlp1 has also been shown to dynamically associate with nuclear pores (Niepel., 2013; Niño et al., 2016), it was nevertheless considered a default component of nucleoplasmic NPCs that prevents the exit of aberrant mRNPs from the nucleus. However, if, as our data demonstrates, the basket is not a default structure of every NPC, what are the consequences for its presumed function as a nuclear gatekeeper. Moreover, if its assembly to a subgroup of pores is linked to its function in quality control and/or mRNA export, one could expect the loss of proteins that have been linked to Mlp1 function to affect its assembly at NPCs. Yet depletion of mRNA maturation factors linked to Mlp1 at the pore such as Yra1, Nab2, or Sac3 (Fischer et al., 2002; Green et al., 2003; Zenklusen et al., 2001) did not affect basket formation at the periphery. Nab2 is thought to mediate the interaction of mRNPs with the nuclear pore by binding directly to the C-terminal region of Mlp1, which is believed to act as the docking site of mRNPs at the basket (Fasken et al., 2008; Green et al., 2003). Nab2, but not Mlp1 is essential for mRNA export, however, over-expression of the C-terminal region of Mlp1 required for Nab2 and hence the mRNP interaction, leads to the accumulation of poly(A) RNA in the nucleus, possibly by sequestering mRNPs away from the pore, as it localizes throughout the nucleoplasm (Fasken et al., 2008; Green et al., 2003). Similarly, depletion of Yra1, which dissociates from mRNPs upon its ubiquitination after pore-docking in a suggested re-arrangement step, and which was shown to accumulate with Mlp1, Nab2, and poly(A) mRNA in nuclear granules upon heat-shock (Carmody et al., 2010), does not affect assembly of Mlp1 at the nuclear periphery. Neither does loss of Sac3, a TREX-2 component involved in TREX-2 association at the pore as well as assisting the recruitment of genes associated with the transcriptional coactivator complex SAGA to the NPC (Jani et al., 2009). The lack of effect on Mlp1-localization at the periphery upon loss of these proteins indicates that basket assembly is separated from the basket’s role in mRNP binding to the NPC and mRNA export. This is further underlined by the sustained presence of Mlp1 at the periphery in the absence of Mex67.

On the other hand, production of mRNA per se appears to be a requirement for the basket assembly as shut-down or depletion of RNA Polymerase II leads to an increase in nucleoplasmic Mlp1 and formation of a distinct Mlp1-granule at the periphery. More specifically, 3’end processing and polyadenylation of transcripts seem to be crucial for basket assembly as loss of distinct factors of the 3’end processing and polyadenylation machinery interferes with Mlp1 localization at nucleoplasmic pores. Loss of the Cleavage Factor IA (CFIA) complex components Rna14 and Rna15 resulted in the assembly of nucleolar pores similar to the phenotype observed upon depletion of the ribosome biogenesis factor Enp1 and nuclear RNA exosome component Csl4. In both *csl4-ph* and *enp1-1* cells, poly(A) pre-rRNA and snoRNA transcripts were previously found to accumulate and sequester the poly(A)-binding protein (PABP) Nab2 into the nucleolus together with poly(A) mRNA, a phenotype that was partially rescued by overexpression of the PABP Pab1, while Nab2 remained nucleolar (Aguilar et al., 2020). Nucleolar poly(A)RNA accumulation has also been observed in yeast strains carrying the temperature-sensitive lethal alleles *rna14-1* and *rna15-2* under restrictive temperature (Carneiro et al., 2007). It is likely, as poly(A) mRNA levels are severely reduced in the absence of Rna14 and Rna15, that Nab2, together with the remaining mRNA maturation machinery and RNAs, is also sequestered into the nucleolus in these cells. However, why the nucleolar sequestration of Nab2 would lead to an increase in basket assembly on NPCs along the nucleolar periphery, and whether these pores participate in any nuclear RNA export, remains unclear (Aguilar et al., 2020; Carneiro et al., 2007), especially since the interaction between Nab2 and Mlp1 is not required for basket assembly at pores.

Pab1 is also believed to promote mRNA export by interacting with different nuclear export factors as well as protecting transcripts from nuclear decay (Schmid et al., 2015; Baejen et al., 2014); however, while no interaction of Pab1 with Mlp1 has been reported to date, its depletion impacted basket assembly with a concomitant increase in nucleoplasmic Mlp1 levels, formation of a large Mlp1-focus at the periphery and ectopic basket formation. Pab1 interacts with components of the CFIA, particularly Rna15 (Amrani et al., 1997), and binds upstream of the poly(A)-site (Baejen et al., 2014). Moreover, Pab1 was shown to recruit the poly(A)-dependent nuclease PAN for control of poly(A)-tail length and 3’end processing, and deletion of PAB1 in suppressor strains resulted in exosome-dependent transcript retention at sites of transcription (Amrani et al., 1997; Brown et al., 1996; Dunn et al., 2005; Minvielle-Sebastia et al., 1997). It is therefore possible that it is the release of a fully 3’ end processed and polyadenylated transcript from the site of transcription that mediates basket assembly, as basket formation is also decreased in the absence of the Poly(A) Polymerase Pap1. While transcription, transcript processing, and release appear to be mediators of basket formation at NPCs, splicing is not. Among the various spliceosome factors depleted in our study only loss of Prp5 affected basket assembly. In addition to its role in pre-spliceosome assembly, Prp5 was shown to regulate transcription initiation/elongation and proofreading in cooperation with SAGA, and the mutant *prp5–1* exhibited decreased RNA Pol II recruitment to intron-containing genes (Shao et al., 2020). The latter suggests that impairment of basket formation upon Prp5 depletion is likely caused by a decrease in mRNA production but not inhibition of splicing, and that splicing itself is not required for basket assembly at NPCs.

Of the proteins involved in later steps of nuclear RNA metabolism only depletion of Pml39 impaired basket assembly with an observed phenotype similar to that of Pab1 mutants. Pml39 and Mlp1 have both been linked to the surveillance of intron-containing mRNAs as the deletion of either led to a leakage of unspliced transcripts to the cytoplasm, whereas their codeletion did not exacerbate this phenotype suggesting a role along the same pathway (Bonnet et al., 2015). Moreover, either protein is required for the association of the other with nuclear pores suggesting that Mlp1 and Pml39 might stabilize each other at the periphery ( Palancade et al, 2005). Hence, only Pml39 connects formation of the basket with its function, which, while still unclear, appears to assemble a functionally specialized interactome.

### A functionally specialized Mlp1 and basket

Baskets do not assemble on every pore along the nucleoplasmic periphery suggesting that having a basket is not the default state for a pore but rather depends on nuclear events along the mRNA metabolism pathway. While their distribution seems to follow a stochastic pattern, baskets do associate with a selected protein and RNA interactome. Overall, our data suggests that both types of pores – basket-containing and basket-less – are export competent consistent with the fact that Mlp1/2 are not essential for export per se (Galy et al., 2004; Green et al., 2003; Powrie et al., 2011). The observation that not all NPCs assemble a basket raises the question of what initiates the assembly at only some pores, beyond the release of correctly processed and polyadenylated mRNAs from their sites of transcription. As basket assembly requires mRNA, however, and a small subset of RBPs, it is possible that only a specific subset of mRNPs triggers basket formation and preferential transit through basket-containing pores. Both Pml39 and the TREX-2 complex were both highly enriched with Mlp1/basket-containing pores yet absent from basket-less NPCs. Mlp1/the nuclear basket and Pml39 have both been linked specifically to surveillance and retention of intron-containing transcripts at the pore. TREX-2, however, has mostly been implicated in the recruitment of transcribing genes to the NPC in connection with SAGA (Jani et al., 2009; Jani et al., 2014). SAGA-regulated genes represent only ~10% of genes in yeast, many of them regulated or induced by different stresses or changes in environmental conditions (Holstege et al., 1998) and no such conditions were induced in our experiments, suggesting that TREX-2 may serve additional functions at basket-containing pores in conjunction with Mlp1. The observation that a certain Mlp1/basket interactome, including TREX-2, is still found at nucleolar pores upon Enp1 depletion might suggest that basket assembly is not linked to an association of actively transcribing genes at the periphery as it is improbable that chromatin will move to the nucleolus under these conditions.

Overall, these observations might further suggest that the location of basket formation at NPCs depends on a balance of Mlp1 concentration, mRNA production and release, and specific RBPs (Figure 8A, left) rather than subnuclear compartments. It has been believed that the nucleolus may form a barrier and that Mlp1 cannot enter the nucleolar space; however, free diffusing Mlp1 is not excluded from the nucleolus and it is the multimerization of Mlp1 during the formation of a basket or granule that causes its exclusion from the nucleolus, as upon the formation of ectopic or nucleolar baskets the nucleolus appears to either form an invagination or rounds up and internalizes, respectively, even forming more than one nucleolus in certain instances (Figure 8A, right). This, as well as the assembly of a basket on only some pores, may, on the other hand, suggest that basket assembly is a stochastic process. Upon Pml39, Prp5, Pap1, and Pab1 depletion and, to a lesser extent, upon shut-down of RNA Pol II, we observed an Mlp1-GFP signal remaining at the nuclear periphery. This observed signal is significantly weaker than that of a basket in wild type cells and, in comparison, continuous along the nuclear rim, including the nucleolar periphery. This may be the consequence of a number of Mlp1 molecules binding to NPCs but below the required stoichiometry of fully formed baskets (Figure 8A, middle). This suggests that self-assembly of Mlp1 at the NPC is not enough to form a functional basket but instead requires other processes along the mRNA maturation pathway. One such process could be the pre-association of Mlp1 with mRNPs in the nucleoplasm as Mlp1 has been found co-isolating with mRNPs (Oeffinger et al., 2007) and has a free-diffusing nucleoplasmic pool. Reaching the pore as part of an mRNP could then contribute to basket assembly, either via stabilization or recruitment of more Mlp1, a notion that would also be in line with assembly of nucleolar pores in *enp1-1* and *csl4-ph* mutants.

**Figure 8.**
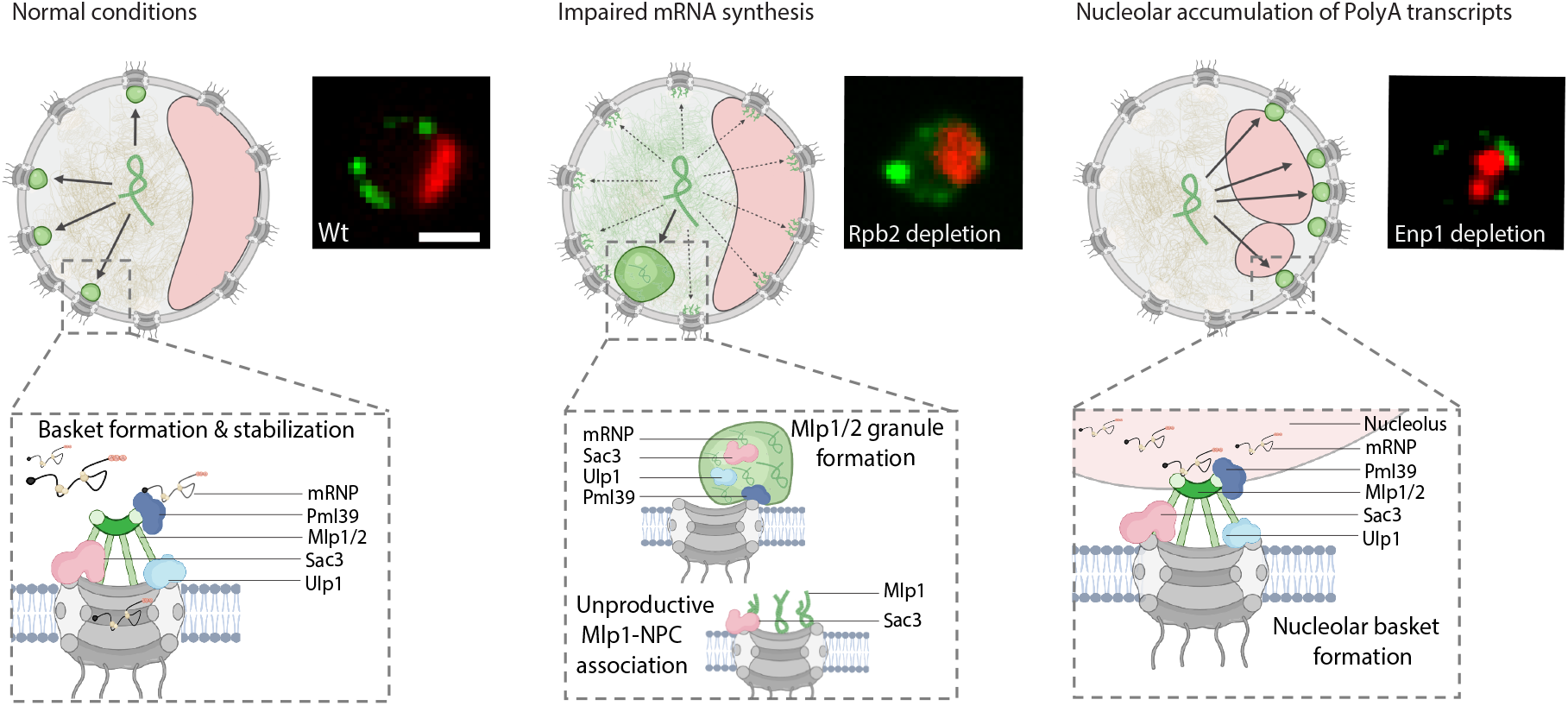
mRNA metabolism drives basket assembly. Cartoons of yeast nuclei illustrating different models of Mlp1-assemblies: either assembly of productive nuclear baskets (left), unproductive Mlp1-NPC binding and Mlp1 granules (middle), and nucleolar baskets (right). Nucleoli are shown as red crescents, Mlp1 as green ampersands. Fluorescent images representing the three different types of Mlp1 assemblies (green) and their localization with respect to the nucleolus marked in red by Gar1-tdTomato are shown on the right. See text for details.

Alternatively, Mlp1 may need to reach a critical concentration at the NPC to form a basket. Our observations show that Mlp1 is an aggregation-prone protein as inhibition of RNA Pol II transcription as well as Pap1 and Pab1 depletion cause granule formation at the nuclear periphery similar to that observed upon loss of Nup60. As under these conditions the level of nucleoplasmic Mlp1 is increased, and, moreover, Mlp1-granule formation seems to scale with the severity of the loss of Mlp1 from the periphery (Figure 8A, middle), basket formation may be dependent on a fine balance of Mlp1 concentration and mRNAs/mRNPs in flux towards the periphery. Mlp1 granule formation, or aggregation, is then prevented by its contact with mRNPs. However, this capacity for aggregation could have been retained by cells either to rapidly assemble granules for sequestration of specific mRNPs and/or to remove baskets and basket-mediated processes from the pore under stress conditions such as heat shock.

### The role of the basket in selective transport

Previous data has suggested that the basket might serve as a quality control platform ensuring that partially or unspliced RNAs do not exit the nucleus. However, the literature concerning the role of the basket in the retention of intron-containing RNAs is conflicting as, in general, the majority of intron-containing mRNAs in yeast are short, while observations of leakage and retention of intron-containing transcripts in both cases are based on a ~3kb reporter transcript that is inefficiently spliced (Palancade et al., 2005; Bonnet et al., 2015; Galy et al., 2004). We observed that while most mRNAs preferentially associated with Mlp1/basket-containing NPCs, there was a clear underrepresentation of short mRNAs in these samples, which include the majority of intron-containing transcripts. Moreover, recent data in human cells did not observe an increase in the leakage of endogenous pre-mRNAs upon TPR depletion (Aksenova et al., 2020; Lee et al., 2020; Zuckerman et al., 2020), different to what was found previously when using inefficiently spliced reporter constructs (Rajanala et al., 2012). These studies also showed that transcripts whose export depends on TPR—and human TREX-2—were in general short, intron-poor or intron-less (Aksenova et al., 2020; Lee et al., 2020; Zuckerman et al., 2020). However, it is important to note that while one of these features is low number of introns, it is not—as previously thought—the mere presence or absence of introns, suggesting the basket may not be primarily a designated gatekeeper for unspliced pre-mRNAs. Yet, it is possible that the function of a basket in pre-mRNA retention becomes more pronounced in a situation where splicing is inefficient.

Overall, Mlp1/TPR-dependent transcripts may have common features, which would point towards a role of the basket in selective transport. However, while our data also points towards such a selectivity model, it is, at least for Mlp1 in budding yeast, unclear what this selectivity involves. As was suggested by data in human cells, features such as short transcript length could indicate that some mRNAs require the nuclear basket to be efficiently exported; short mRNAs and those that have undergone only one to two splicing events may represent less complex mRNPs lacking components to facilitate efficient binding to and transport through the pore. If we consider mRNP scanning of the nuclear periphery part of a probabilistic process towards successful binding to the pore through specific RBP:NPC interactions, it is possible that shorter transcripts may require the basket to facilitate such a binding event. In line with such a model, recent APEX2 data found that short mRNAs were enriched with NPCs in human cells, possibly due to a lower number of NPC-interacting factors on these transcripts that would mediate efficient translocation (Fazal et al., 2019). Moreover, a relationship between length and number of RBPs has also been suggested by previous *in silico* predictions that indicate basket-mediated mRNA quality control to be a length-dependent mechanism (Soheilypour et al., 2016).

Alternatively, the basket might mediate an increased residence of longer mRNAs at the pore prior to export, creating a kinetic delay for larger mRNPs that require rearrangements and/or quality control. Such a model would be in line with the enrichment of longer mRNAs observed with Mlp1/basket-containing pores, including those containing introns, and would further explain the dependence of the 3kb-intron-leakage/retention reporter on Mlp1/2. In addition, supporting the notion of a remodeling step that facilitates mRNP rearrangements prior to export, previous studies showed that ubiquitination of Yra1 by the E3 ligase Tom1 at the pore promoted its dissociation from mRNPs (Iglesias et al., 2010), whereas deletion of TOM1 resulted in the nuclear retention of poly(A) mRNAs and their localization within nuclear foci together with Nab2 (Duncan et al., 2000).

While the nuclear basket is not a requirement for nuclear mRNA export per se, it nevertheless appears to serve as a facilitator. Either model for the nuclear basket as mediator of selective export kinetics, whether it is to enhance or slow access to the central pore channel for translocation, would also be compatible with different models of basket formation and function. Independent of which of the many proposed functions of the nuclear basket will prevail as its primary role in mRNA transport and mRNA metabolism overall, our study shows that in budding yeast, this function is dedicated to a select subset of nuclear pores. Whether a similar specialization of nuclear pores exist in other organisms and whether pore specialization extends to other functions of the NPC is an exciting topic for future research.

## ACKNOWLEDGEMENTS

We thank S. Adivarahan, P. Raymond, L.C. Aguilar and C. Trahan for discussions and help in the optimization of different experimental approaches. We thank C. Strambio-De-Castillia for generously providing plasmids encoding yEGFP3-NLS-Mlp1 truncations (Niepel et al., 2013); L. Lavis for HaloTag ligands; M. Swaffer, J. Skotheim, and R. Reyes-Lamothe for the yeast strain expressing Halo-NLS. Furthermore, we like to thank D. Faubert at the IRCM Proteomics Platform, the IRCM Molecular Biology Platform, and N. Stifani at the BMM microscopy platform for help. This work was supported by Natural Sciences and Engineering Research Council Discovery Grants to D.Z. (RGPIN-2015-05922) and M.O. (RGPIN-2015-06568), a joint Canadian Institute for Health Research Project grant to D.Z. and M.O. (PJT-425798), the Canadian Foundation for Innovation (D.Z. and M.O.), and the National Institutes of Health to B.M. (R01-GM124120). D.Z. holds a FRSQ Chercheur Boursier Senior.

## MATERIAL AND METHODS

### Yeast tagging and growth

Yeast strains and plasmids used in this study are listed in Tables S2 and S4. Yeast strains are all derived from W303, and epitope-tagged proteins have been C-terminally tagged. Yeast strains were constructed by homologous recombination using a recombination cassette generated by PCR (for primers, see Table S3) with 50 bp homology arms as described in (Bensidoun et al., 2016). Cells were grown in YPD or synthetic complete media lacking the appropriate amino acid to maintain plasmids when appropriate. Unless noted otherwise, cells for RNA-based or protein-based analyzes were isolated, rapidly frozen in liquid nitrogen, and cryo-lysis was performed by solid phase milling in a planetary ball mill (Retsch) producing a fine cell grindate (Oeffinger et al., 2007). All grindates were stored at −80°C until processed either for affinity purification or RNA extractions.

### Mlp1 N-terminal fragment expression

All expression plasmids encoding yEGFP3-NLS-Mlp1 truncations have been generously provided by C. Strambio-De-Castillia (Niepel et al., 2013). Cells transformed with these expression plasmids were grown in the presence of 150 mg/l methionine for live-cell microscopy to reduce the expression level of the GFP-NLS–tagged Mlp1p fragments in target yeast cells.

### Auxin depletion

Yeast cells were grown at 30°C in YPD to an OD_600_ ~0.3. Indole 3 acetic acid (auxin; Sigma Aldrich) was added to a final concentration of 500 μM (Morawska et al, 2008). After 120 min, cells were prepared for live-cell imaging in SD with 2% glucose supplemented with 500 uM of auxin. For western blotting, 20 ml of culture was harvested at an OD_600_ of ~0.6 with or without 500 μm auxin, and protein extraction and western blotting carried out as described above.

### Protein extraction and western blot analysis

For western blot analysis, proteins were extracted from yeast cells was performed as previously described in (Kushnirov et al., 2000) and separated on 10% or 4 – 12% Tris/Glycine SDS PAGE gels. Proteins were transfer to PVDF membranes and detected using either monoclonal anti-GFP (1:1000; Sigma, 11814460001), anti-HA 12CA5 (1:1000; Sigma, 11583816001), or anti-mouse HRP (1:5000; Abcam, ab6728) antibodies. Images were acquired using a ChemiDoc MP Imaging System (Biorad).

### Preparing cells for live-cell imaging

Yeast cells were grown at 30°C in SD with 2% glucose to an OD_600_ ~ 0.4–0.6. For imaging, 100-μl cell suspension was added to a 96-well glass-bottom plate (MGB096-1-2-LG-L; Brooks Life Science Systems) previously coated with concanavalin A (Con A) and concentrated on the bottom of the well by centrifugation. Wells were coated by adding 100 μl of 1 mg/ml Con A (Sigma-Aldrich) for 10 min before unbound Con A is removed and the Con A activated by adding 100 μl of 50 mM CaCl_2_/50 mM MnSO_4_ for 10 min. The solution was then removed, washed once with 100 μl ddH_2_O, and air-dried. To minimize cell motion for SIM imaging, cells were briefly fixed with 70% EtOH for 5 min chilled at −20C, wash with cold 1x PBS before been added to glass-bottom plates coated with ConA.

### Cell labeling with HaloTag ligands

Cells were grown in YPD to an OD_600_ ~0.15 in log phase before incubating with 100 nM of Halo-ligand JF-549 (generously provided by Luke Lavis, Janelia Research Campus) (Ling et al., 2015) for 90 min, followed by 3 quick YPD washes and a 30min wash in YPD with agitation. Finally, cells were washed 3 times in SD with 2% glucose before imaging. For SIM experiments, cells were fixed after the last wash as described above.

### Image acquisition

Unless mentioned otherwise in the text, images were acquired on a spinning disk confocal microscope (Observer SD; Carl Zeiss) using a 100×/1.43 NA objective (Carl Zeiss), 488-nm (100 mW), and 561-nm (40 mW) excitation laser lines, and Semrock single bandpass filters for GFP (525 nm/50 nm) and RFPs (617 nm/73 nm). Images were captured using an electron-multiplying charge-coupled device camera (Evolve 512; Photometrics) using Zen blue software. For heat-shock experiments, the stage was preheated to 42°C using a Zeiss incubation chamber. Cells were incubated in the chamber for 1h at 42C in SD with 2% glucose.

### Structured illumination microscopy (SIM)

SIM images were acquired with a 63x NA 1.46 oil objective on a Zeiss Elyra PS.1 system equipped with an Andor EMCCD iXon3 DU-885CSO VP461 camera (1004×1002 pixels), and with the following lasers: 50 mW405 nm HR diode, 100 mW 488 nm HR diode, 100 mW 561 nm HR DPSS, 150 mW 642 nm HR diode. Each image was acquired using 3 rotations and a grid size of 42mm for all channels.

### Single molecule imaging using Highly Inclined Laminated Optical (HILO) illumination

Cells were concentrated and mounted for imaging as described above. Images were acquired using Zeiss Elyra PS.1 microscope setup equipped with a 63x NA 1.46 oil objective, an Andor EMCCD iXon3 DU-885CSO VP461 camera (1004×1002 pixels) and with the following lasers: 50 mW405 nm HR diode, 100 mW 488 nm HR diode, 100 mW 561 nm HR DPSS, 150 mW 642 nm HR diode. To minimize the out-of-focus light, time-lapse movies were acquired using Highly Inclined Laminated Optical (HILO) sheet illumination. Image size was cropped to 512×512 pixels, and images acquired at 50 Hz (frames per second). To quantify single molecule diffusion in the nucleoplasm and the nucleolus, images were acquired until most fluorophores were bleached and single molecules became visible to allow single particle tracking. Tracking was done using the Trackmate plugin in ImageJ (Tinevez et al., 2017). To measure the fraction of tracks from single diffusing particles overlapping with the nucleolus, the nucleolar areas were first manually delimited using the ROI manager tool in ImageJ, followed by counting the fraction of time individual single particles overlapped with the nucleolus. 40 tracks for Mlp1-Halo and Halo-NLS, varying from 10 to 40 frames were analyzed.

### Affinity purification and mass spectrometry

#### Affinity purification

Affinity purifications (AP) were performed in triplicate per conditions as previously described (Oeffinger et al., 2007b). In brief, cells were grown to late log phase, frozen by immersion in liquid nitrogen, and mechanically ground using a planetary ball mill (Retsch). For each AP, 1 g of cell powder was thawed in 9 ml of extraction buffer (1X tributyltin, 50 mM NaCl, 1 mM DTT, 0.5 % Triton X-100, 1X protease inhibitor cocktail [4 mg/mL pepstatin A (Sigma), 180 mg/mL PMSF (Sigma)], antifoam B (Sigma, 1:5000), and 40U/mL RNAsin (Promega)), homogenized with a Polytron for 25 s, and cleared by centrifugation at 4,000 g for 5 min. 10mM glutaraldehyde was added for 5 min and samples were gently agitated on ice before the reaction was quenched with Tris-HCl pH8 to a final concentration of 100mM. Lysates were incubated with either IgG (anti-rabbit IgG, Sigma) or GFP-nanobody (expressed from pDZ580-pET28a-GBP and purified as described in-conjugated magnetic beads (Dynabeads M-280) for 30 min (Rothbauer et al., 2008). After removal of the super-natant, beads were extensively washed in extraction buffer, then detergents removed by washing the bead-bound complexes in 0.1 M NH_4_OAc/0.1 mM MgCl_2_ before a final was and resuspension in 50μl of 20-mM Tris-HCl, pH 8.0. Isolated proteins were digested on-bead at 37°C with 1 μg trypsin (Pierce Trypsin Protease, MS Grade) for 16 h (Trahan et al., 2016). The digestion was stopped by adding formic acid to a final concentration of 2%.

For differential APs, the flow-through was incubated with GFP-nanobody-conjugated magnetic beads for 30 min and the beads then treated as describe above.

#### Peptide preparation for injection in mass spectrometer

Tryptic peptides were cleaned using C18 ZipTips as per supplier recommendations (Milli-pore). Samples were injected to near saturation of the signal, while an equivalent volume of their respective negative controls were injected. Liquid chromatography was performed using a PicoFrit fused silica capillary column (15 cm × 75 μm i.d; New Objective, Woburn, MA, USA), self-packed with C-18 reverse-phase resin (Jupiter 5 um particles, 300 Å pore size; Phenomenex, Torrance, CA, USA) using a high-pressure packing cell on the Easy-nLC II system (Proxeon Biosystems, Odense, Denmark) and coupled to an Orbitrap Fusion™ Tribrid™ Mass Spectrometer equipped with a Proxeon nanoelectrospray Flex ion source. 0.2% formic acid (Solvent A) and 100% acetonitrile/0.2% formic acid (Solvent B) were used for chromatography and peptides were loaded on-column at a flowrate of 600 nl/min and eluted with a three-slope gradient at a flowrate of 250 nl/min. Solvent B was first increased from 2 to 25% over 20 min, then from 25 to 45% over 40 min, and finally from 45 to 80% B over 10 min.

#### Protein identification

The peak list files were generated with Proteome Discoverer the following as described in (Aguilar et al, 2020). Protein database searching was performed with Mascot 2.5 (Matrix Science) against the NCBI - *S. cerevisiae* protein database (20160802). The mass tolerances for precursor and fragment ions were set to 10 ppm and 0.6 Da, respectively. Trypsin was used as the enzyme allowing for up to 1 missed cleavage. Cysteine carbamidomethylation was specified as a fixed modification, and methionine oxidation as variable modifications. Data analysis was performed using Scaffold (version 4.8.4).

#### Mass spectrometry data analysis

Protein and peptide identification thresholds in Scaffold™ was set to 95% which resulted in decoy false discovery rate of 6%. Exclusive spectrum counts (ESC) were used for semi-quantification of protein preys, and mass spectrometry results were analyzed as previously described in (Scott et al., 2017). Briefly, only Exclusive Spectral Counts (ESCs) above background detected in controls were retained. In silico digestion using MS digest (http://prospector.ucsf.edu) was performed for each protein to take protein size and predicted cleavage sites into accounts. Values were normalized against the average values of the proteins associated with the bait proteins in the different APs: Mlp2 for Mlp1-PrA and Y complex nucleoporins (Nup84, Nup85, Nup120 Nup145C) for Nup133-GFP. This allowed normalization of the data sets against proteins with a similar size, stoichiometry, and segregation behavior without the need to consider the none-pore associated fraction of bait proteins. Data are available via ProteomeXchange with identifier PXD027872.

Project Name: Nuclear pore complexes interactome dissection

Project accession: PXD027872

Project DOI: 10.6019/PXD027872

**Reviewer account details:**

Username

Password: gyCzPdNT

### Affinity purification RNA-seq

#### Poly(A)-RNA preparation and sequencing

RNA Affinity purification was performed using 1 g of cell powder per triplicate as described above but without crosslinking. Lysates were incubated with either IgG- or GFP-nanobody-conjugated magnetic beads for 30 min. Beads were washed extensively (8 times) in extraction buffer before being resuspended in 1 ml of Trizol (Invitrogen, 15596026) and vortexed vigorously. The total poly(A) library was generated by RNA extraction using 100 mg of cryo-ground cell powder thawed into Trizol in triplicates. RNA was then extracted using the Direct-zol Miniprep Kit (Zymo research, R2050) and Dnase treatment was performed on-column according to manufacturer’s instructions. Samples were resuspended in 30 μl ultra-pure water (Invitrogen, 10977023), and the quality of RNA was assessed by Qbit and Bioanalyzer chip. RNA extracts were Poly(A)-RNA enriched via a Poly(A) mRNA magnetic Isolation Module oligo-dT (NEBNext), and cDNA libraries were prepared using the Kapa RNA HyperPrep Kit (96 rxns, Roche) and TruSeq DNA UDI 96 indexes (Illumina). RNA-sequencing was performed using Nocaseq6000 flowcell S2 PE50.

#### RNA sequencing analysis

RNA-seq reads were trimmed using Trimmomatic (version 0.38.0) to quality trim and remove adapters. Trimmed reads were mapped to the *S. cerevisiae* genome (GCF_000146045.2_R64) using STAR (version 2.5.2a) using default parameters. Genes were counted using htseq-count (version 0.11.3) with parameters -m intersection-nonempty -s yes -r pos.

Differential enrichment analysis was performed using DESeq2 (version 1.28.1) using the Wald test. All five dAPs (Basket^plus^, Basket^minus^, ‘All pores’, IgG, and GFP alone) were compared against the total poly(A) library representative of the total transcriptome to generate log2 fold change values for each transcript. Transcripts with an absolute log2 fold change value > 1 and p < 0.05 were considered significantly differentially enriched. Transcripts that were significantly enriched in the IgG and GFP libraries were removed from further analysis from the Basket^plus^ (IgG), Basket^minus^ (GFP), and ‘All pores’ (GFP) results according to purification type. The R library UpSetR was used to generate upset plots and determined overlapping sets of enriched and depleted genes across libraries. The R library clusterProfiler was used to perform gene set enrichment analysis using Gene Ontology terms for each dAP library. To determine whether enriched transcripts correlated with other expression features, we parsed the *S. cerevisiae* GTF gene annotation file (GCF_000146045.2_R64) to determine the presence of introns and transcript length, and used poly(A) tail lengths as determined previously (Tudek et al., 2021). To normalize counts to compare average transcript expression across libraries, the variance stabilized transformation as implemented in the DESeq2 function vst() was used. T tests were performed using the t.test() function, wilcoxon rank sum tests were performed using the wilcox.test() function, and linear models were generated using the lm() function. All analysis code is available at github.com/montpetitlab/Bensidoun_et_al_2021.

---

**Figure S1. A**. Movies of Mlp1-Halo and Halo-NLS diffusion acquired at 20-ms intervals. For Mlp1, early timepoints of the movie show proteins with steady state localization at the nuclear periphery that bleach progressively during image acquisition. **B.** Individual frames from movie tracking Halo-NLS JF646. White arrows show Halo-NLS JF646 in each frame and the dashed circle presents the nucleolar area. MAX shows the maximum intensity projection of all frames with the path highlighted with white crosses. Halo-NLS JF646 is shown in red, nucleolus (Gar1- GFP) in green. (Scale bar =2μm).

**Figure S2: Mlp1 granule formation in *rpb1-1* cells at non-permissive temperature.** Image of Mlp1-GFP distribution in *rpb1-1* cells shows the formation of a single bright Mlp1-GFP granule at 37°C and line scan intensity plot measuring of Mlp1-GFP (green) intensity. Nup188-tdTomato signal is shown in red. histogram on the right shows the quantification of cells that exhibit Mlp1- GFP granule formation in wild type and *rpb1-1* cells at 25°C and 37°C.

**Figure S3. A.** Mlp1-GFP (green) signal is redistributed to the nucleolar periphery in Csl4^AID-HA^ cells upon auxin treatment (120min). Nucleolus labeled by Gar1-tdTomato (red). **B.** Enp1^AID-HA^ cells display fragmented and internalized spherical nucleoli upon Mlp1-GFP relocalization to the nucleolar periphery upon auxin treatment for 120 min.

**Figure S4. A.** Mlp1-GFP distribution was monitored with respect to the nucleolus in cells where baskets were destabilized upon the addition of auxin and upon heat shock. Graphs represent the overlaps of Mlp1-GFP and Gar1-tdTomato signals (green and red curve respectively). White arrows show Mlp1-GFP signals remaining at the periphery, including in the nucleolar area after basket destabilization. (Scale bar =2μm). **B.** Western blot of total cell lysates from AID-HA-tagged components along the mRNA maturation pathway and NPC associated factors strains at different time points upon addition of auxin in Mlp1-GFP cells. AID-HA tagged proteins and Mlp1-GFP were detected using anti-HA and anti-GFP antibodies, respectively.

**Figure S5. A.** SIM co-localization analysis of cells double tagged for Nup188-tdTomato (red) and select GFP tagged proteins (green). **B.** SIM co-localization analysis of cells double tagged for Mlp1-GFP (green) and select Halo-tagged proteins. (Scale bar =2μm)

**Figure S6. A.** Illustration of the NPCs affinity purifications. All pores were affinity purified via GFP-nanobodies-conjugated beads using yeast cell extracts from an Mlp1-PrA/Nup133-GFP double-tagged strain. Separation of basket-containing and basket-less pores was achieved from an Mlp1-PrA/Nup133-GFP double-tagged strain using a differential affinity purification approach, whereby Mlp1-PrA associated complexed (Basket^plus^) were isolated via IgG-conjugated beads, followed by the isolation of Nup133-GFP associated complexed (Basket^minus^) via GFP-nanobodies from the flow-through. Nup133-GFP associated complexed (Basket^minus^) were also isolated from *Δ mlp1/2/*Nup133-GFP cells. **B.** Histograms showing normalized spectral counts of all proteins co-purified with the different pore APs relative to proteins identified in “All NPCs”. Proteins were grouped according to their cellular functions. **C.** All NPCs, Mlp1-PrA (Basket^plus^), and Nup133-GFP (Basket^minus^) associated complexes were isolated from Mlp1-PrA/Nup133- GFP, *Δ mlp1/2/* Nup133-GFP cells and Mlp1-PrA/Nup133-GFP/Enp1^AID-HA^ cells upon auxin treatment. Proteins were included if identified as a high confidence interaction (i.e., at least five exclusive spectrum counts (ESC) in two biological replicates) and grouped according to their cellular functions. Each protein is presented as log2(FC) over proteins identified in ‘All NPCs’.

**Figure S7. Analysis of transcript features across differential affinity purification samples A.** Histograms representing transcript enrichment in Basket^plus^’, ‘Basket^minus^’ and ‘all pores’ from differential APs over mean transcript expression and intron content in the poly(A) library (grey). **B.** Histograms representing transcript enrichment in Basket^plus^’, ‘Basket^minus^’ and ‘all pores’ from differential APs in relation to transcript length observed in poly(A) library (grey). **BC** Histograms representing transcript enrichment in Basket^plus^’, ‘Basket^minus^’ and ‘all pores’ from differential APs in relation to poly(A) tail length observed in poly(A) library (grey).

