## Supplementary figures and images for "Nuclear mRNA metabolism drives selective basket assembly on a subset of nuclear pores in budding yeast"

### Figure S1

A

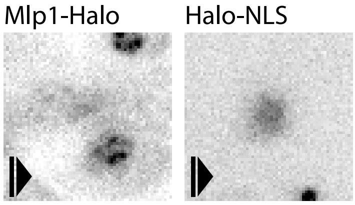


B

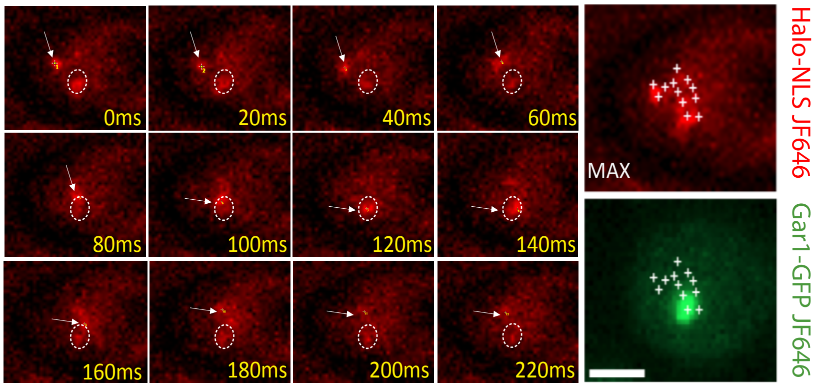


Bensidoun et al. Figure S1

### Figure S2

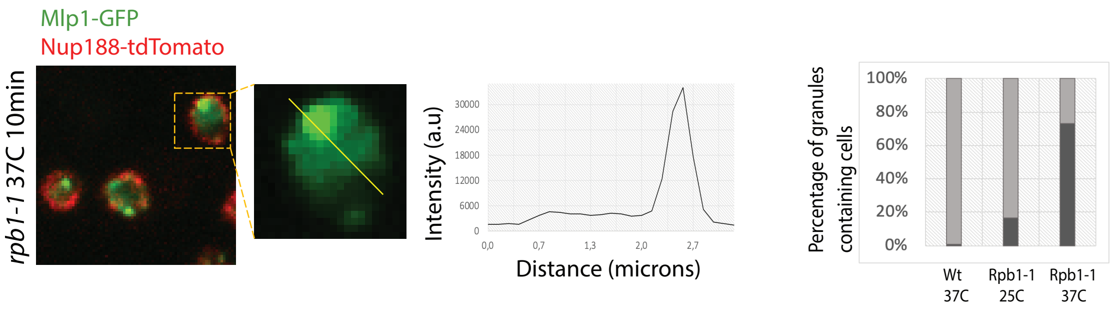


Bensidoun et al. Figure S2

### Figure S3

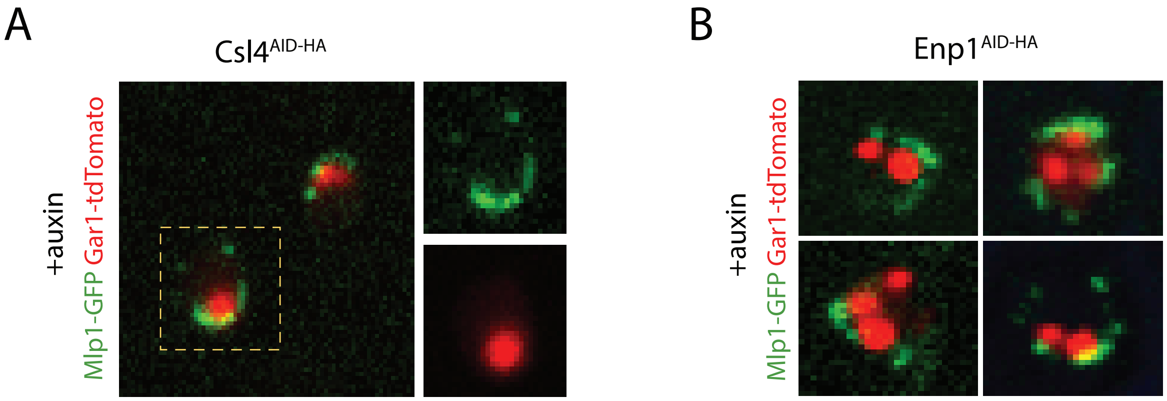


Bensidoun et al. Figure S3

### Figure S4

A

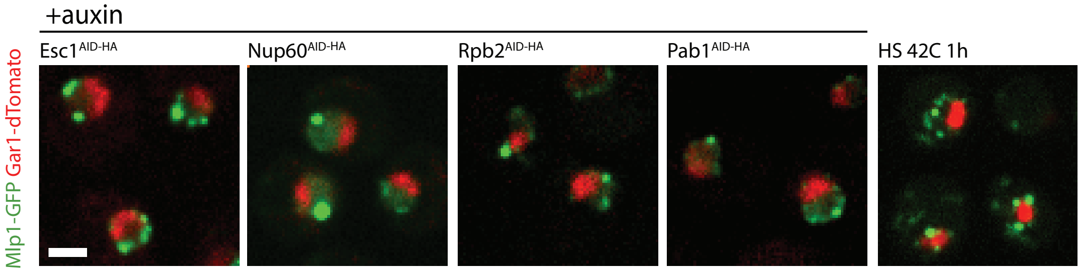


B

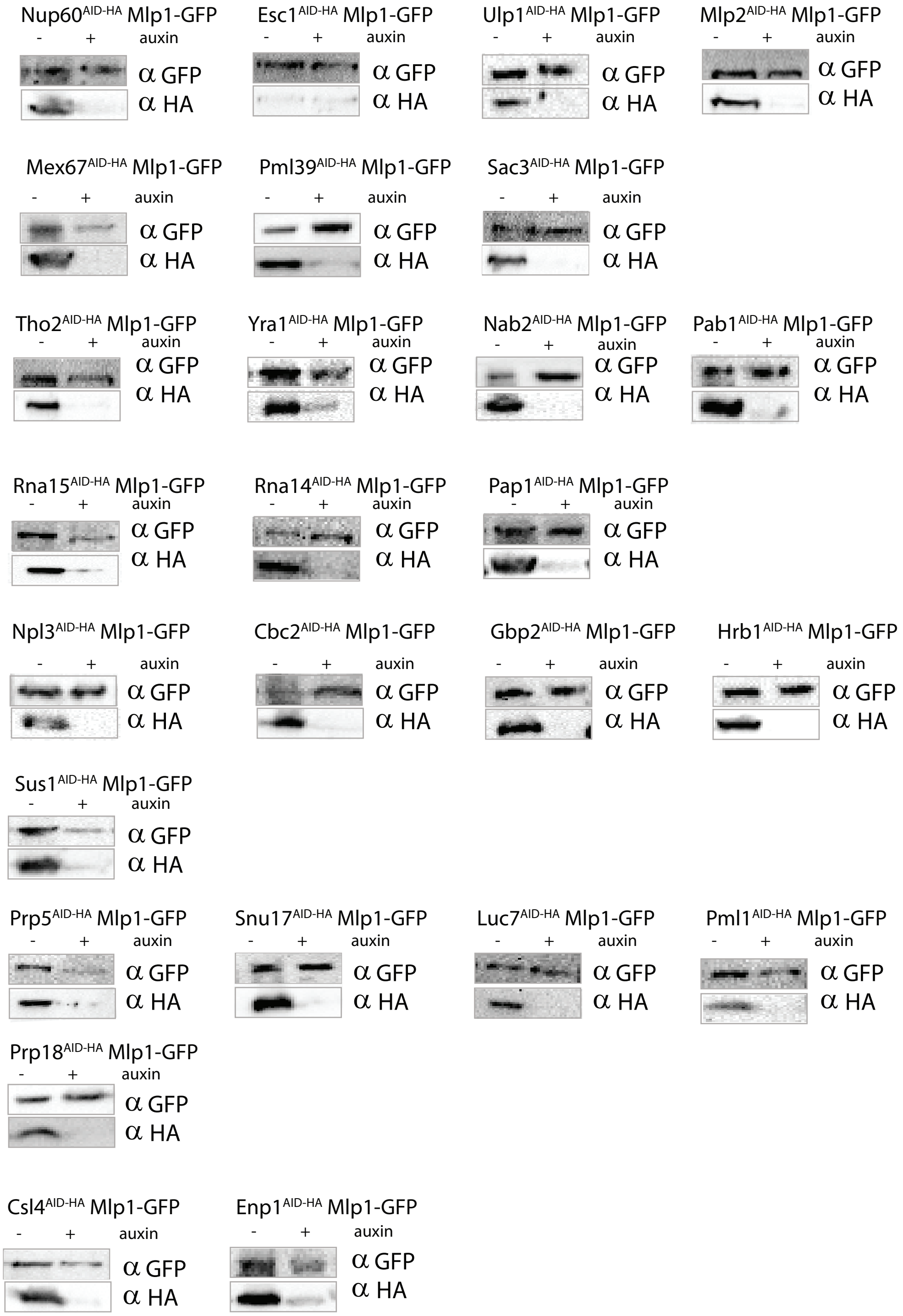


Bensidoun et al. Figure S4

### Figure S5

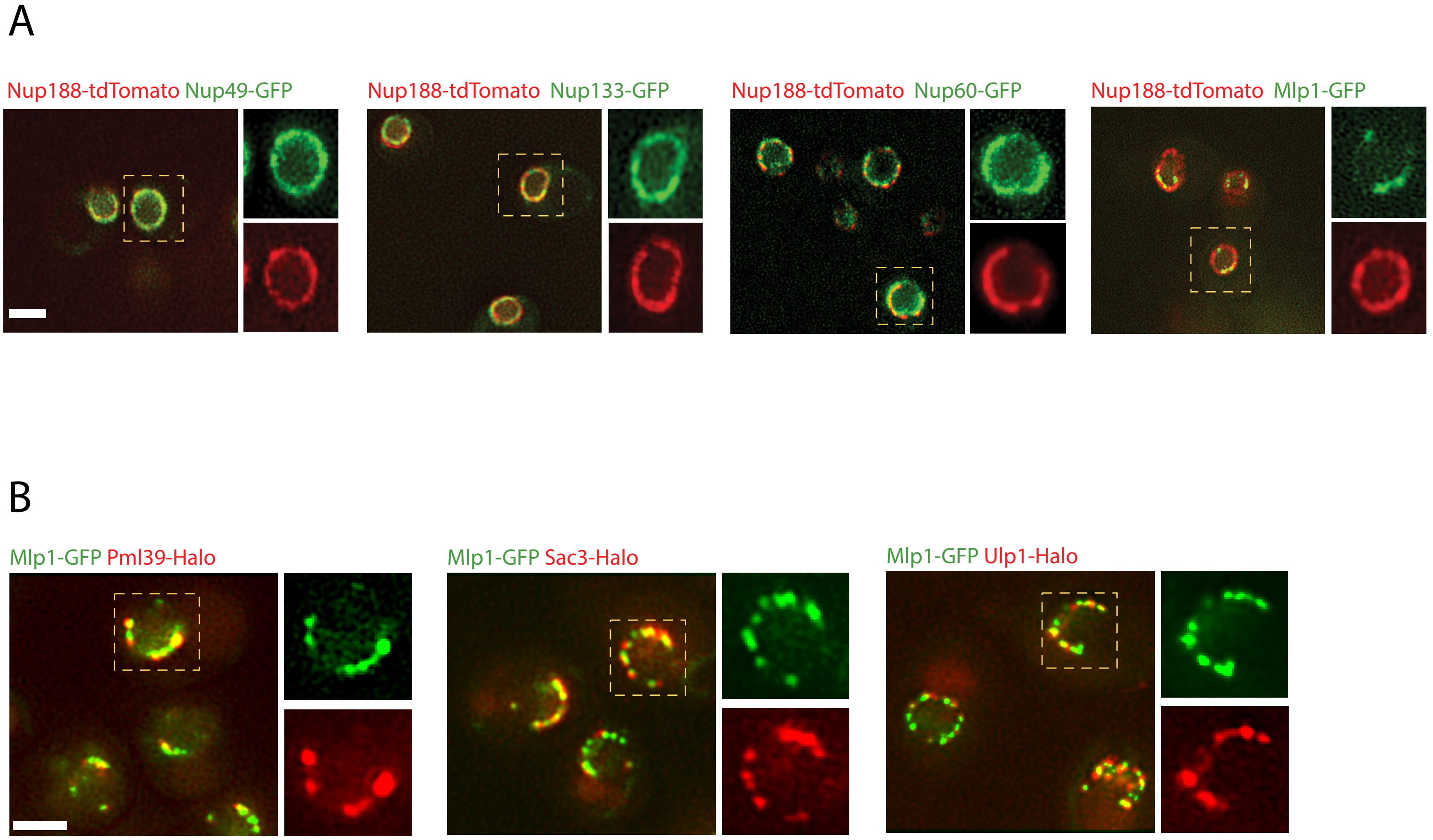


Bensidoun et al. Figure S5

### Figure S6A

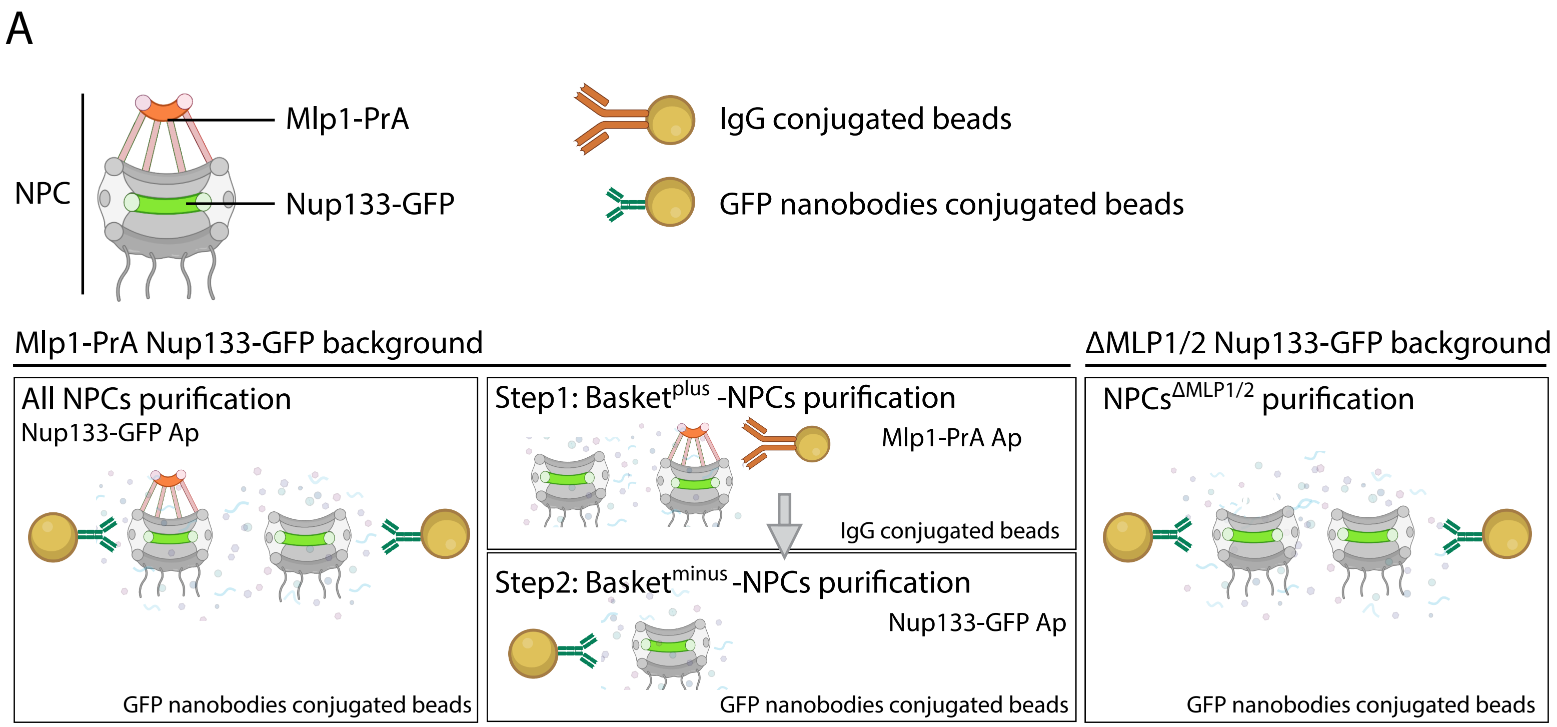


Bensidoun et al. Figure S6A

### Figure S6B

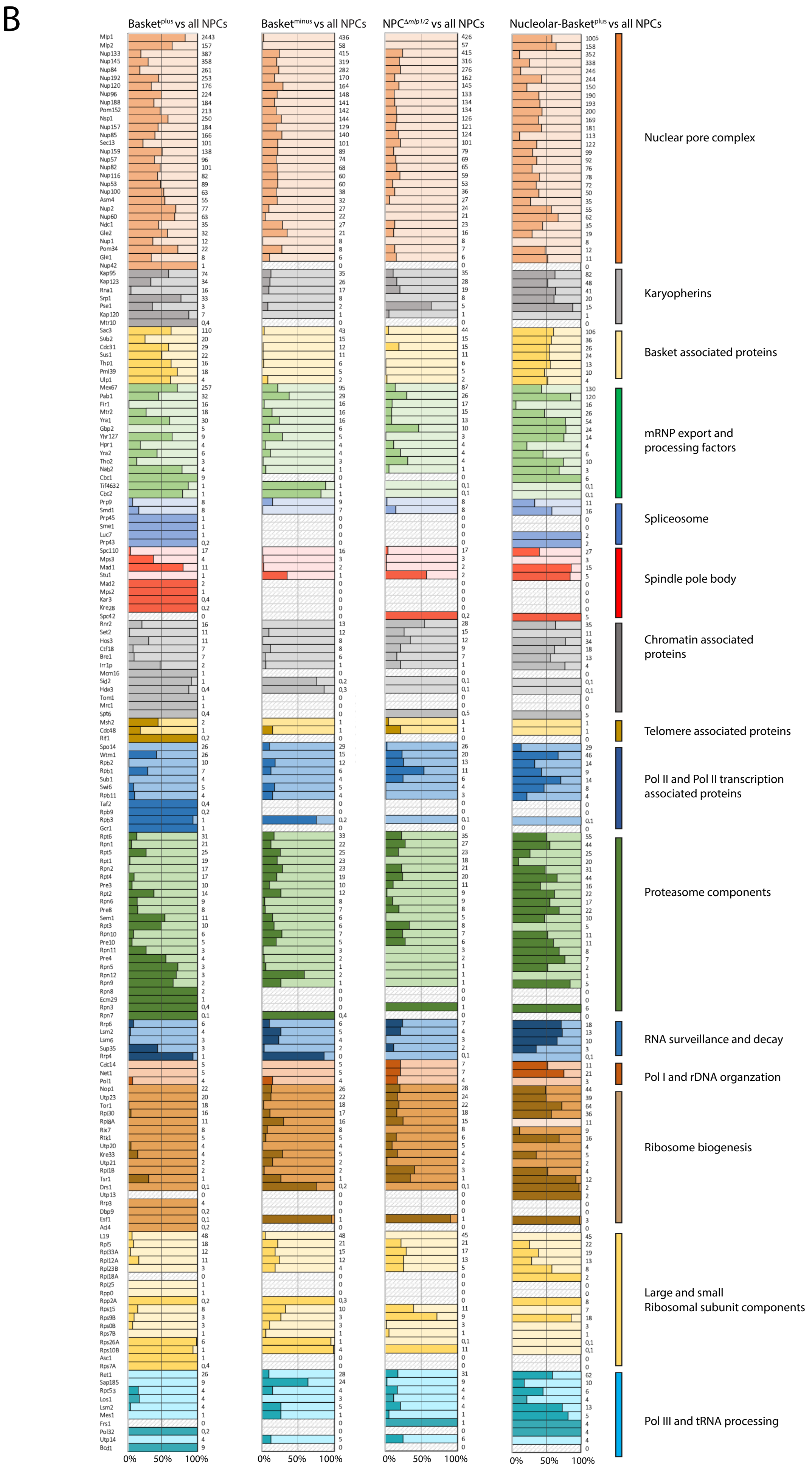


Bensidoun et al. Figure S6B

### Figure S6C

C

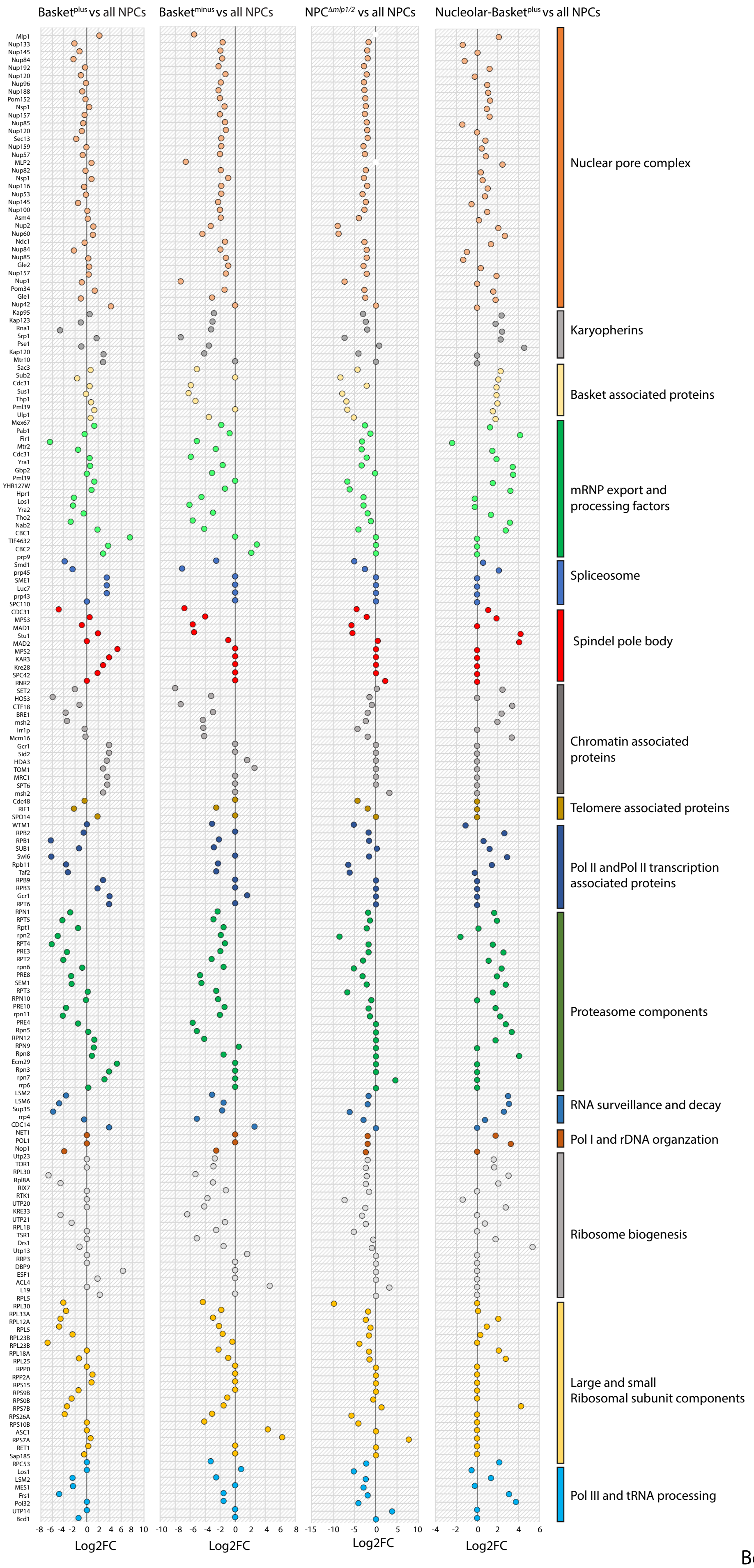


Bensidoun et al. Figure S6C
