## Supplementary material for "Nuclear mRNA metabolism drives selective basket assembly on a subset of nuclear pores in budding yeast": Figure S7

A

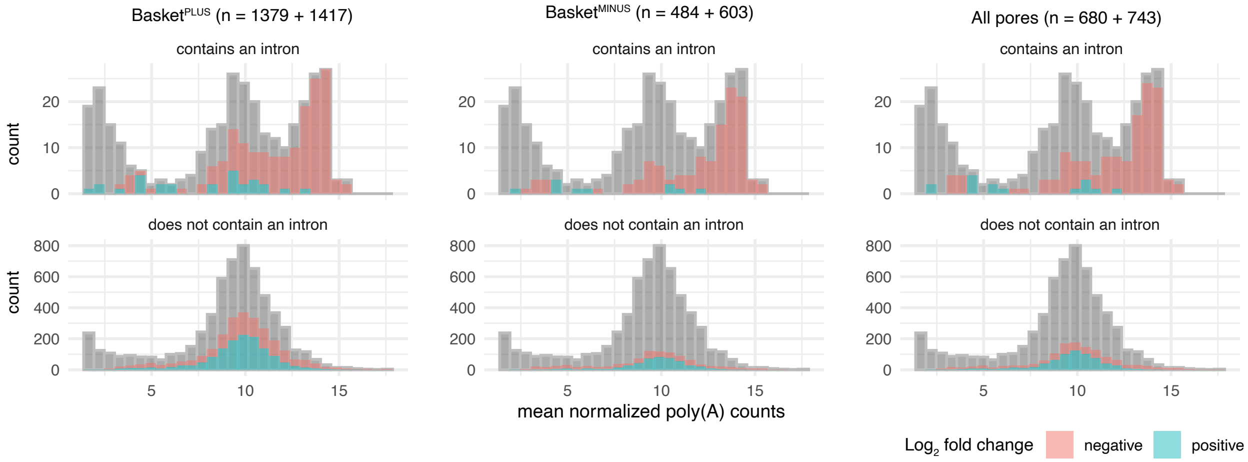


| introns | estimate | statistic | p.value | method |
|---|---|---|---|---|
| no | 0.3361296 | 3.026683 | 0.002517563 | Welch Two Sample t–test |
| yes | 2.4119109 | 2.449554 | 0.022167063 | Welch Two Sample t–test |

| introns | estimate | statistic | p.value | method |
|---|---|---|---|---|
| no | −0.4145561 | −2.483224 | 0.013227581 | Welch Two Sample t–test |
| yes | 5.0240316 | 4.383841 | 0.001329651 | Welch Two Sample t–test |

| introns | estimate | statistic | p.value | method |
|---|---|---|---|---|
| no | 0.1437639 | 1.023376 | 3.063868e−01 | Welch Two Sample t–test |
| yes | 5.0152545 | 5.444374 | 4.962584e−05 | Welch Two Sample t–test |

B

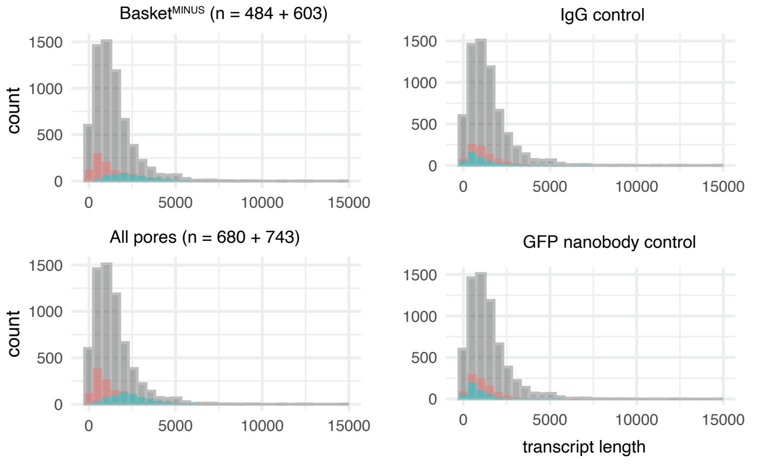


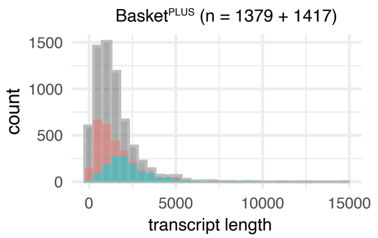


| statistic | p.value | method | set |
|---|---|---|---|
| 118435.0 | 2.168392e−06 | wilcox.test() | IgG ctrl |
| 151490.0 | 3.931193e−14 | wilcox.test() | GFP nanobody ctrl |
| 281372.5 | 2.429320e−234 | wilcox.test() | basket |
| 23500.5 | 1.554581e−125 | wilcox.test() | basketless |
| 35172.5 | 2.711172e−174 | wilcox.test() | all pore |

Log$_2$ fold change negative positive

C

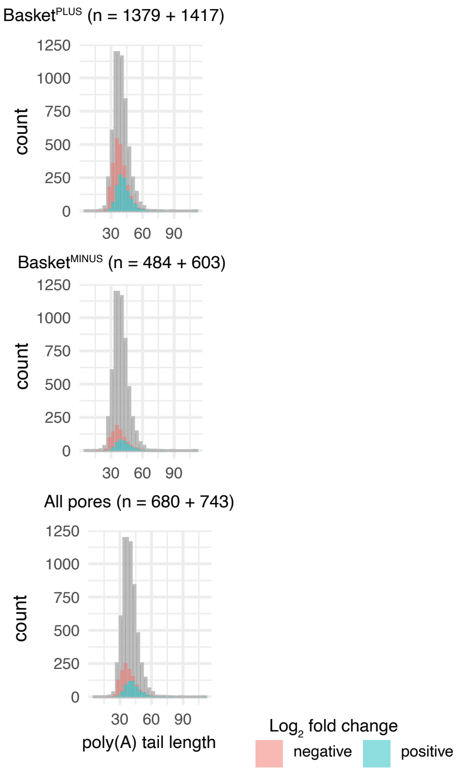


Bensidoun et al. Figure S7
