## Supplementary material for "Nuclear mRNA metabolism drives selective basket assembly on a subset of nuclear pores in budding yeast": Table S1

**Table S1.AID screen candidates and phenotypes**

**All strains generated in this study are constructed in W303 background** *MAT a/MATα (leu2-3,112 trp1-1 can1-100 ura3-1 ade2-1 his3-11,15)*

| Protein | Function | Phenotype upon depletion |
|---|---|---|
| Rpb2 | RNA Pol II subunit | Basket destabilization and Mlp1 granule formation |
| Rpa135 | RNA Pol I subunit | - |
| Enp1 | Small ribosomal subunit export | Basket re-localization at the nucleolar periphery |
| Csl4 | Exosome-non catalytic core component | Basket re-localization at the nucleolar periphery |
| Ulp1 | Ubiquitin-like modifier, Basket associated protein | Basket destabilization |
| Mlp2 | Basket scaffold, Mlp1 homologue | Basket destabilization |
| Nup60 | Basket nucleoporin | Basket destabilization and Mlp1 granule formation |
| Esc1 | Lamin-like protein | Basket destabilization and Mlp1 granule formation |
| Pml39 | Pre-mRNA surveillance, basket associated protein | Basket destabilization |
| Mex67 | Export receptor | - |
| Tho2 | Co-transcriptional mRNP packaging | - |
| Yra1 | TREX subunit ; mRNA export factor | - |
| Sus1 | SAGA and TREX-2 subunit | - |
| Sac3 | TREX-2 scaffold ; mRNA export factor | - |
| Pap1 | Poly(A) polymerase | Basket destabilization and Mlp1 granule formation |
| Nab2 | Poly(A) binding protein | - |
| Pab1 | Poly(A) binding protein | Basket destabilization and Mlp1 granule formation |
| Rna14 | Cleavage and mRNA polyadenylation | Basket re-localization at the nucleolar periphery |
| Rna15 | Cleavage and mRNA polyadenylation | Basket re-localization at the nucleolar periphery |
| Cbc2 | Cap binding complex subunit | - |
| Prp5 | Pre-spliceosome formation; RNA helicase | Basket destabilization |
| Snu17 | Splicing factor; U2 snRNP complex | - |
| Luc7 | Splicing factor; U1 snRNP complex | - |
| Pml1 | Pre-mRNA surveillance | - |
| Npl3 | Nuclear mRNA surveillance | - |
| Gbp2 | Nuclear mRNA surveillance | - |
| Prp18 | Splicing factor; component of snRNP U5 | - |
| Hrb1 | Nuclear mRNA surveillance | - |
