## Supplementary material for "Nuclear mRNA metabolism drives selective basket assembly on a subset of nuclear pores in budding yeast": Table S2

**Table S2. Yeast Strains**

**All strains generated in this study are constructed in W303 background** *MATa/MATα (leu2-3,112 trp1-1 can1-100 ura3-1 ade2-1 his3-11,15)*

| Strain | Description | Reference |
|---|---|---|
| 6772 | Mlp1-GFP, Gar1-tdTomato | This study |
| 6773 | Delta Mlp1/Mlp2, Gar1-tdTomato, Nterm2-GFP | This study. Nterm2 expression Plasmid generated in Niepel et al., 2005 |
| 6775 | Nup188-tdTomato, Nterm2-GFP | This study. Nterm2 expression Plasmid generated in Niepel et al., 2005 |
| 6711 | Mlp1-Halo, Gar1-GFP, Pdr5 delta | This study |
| 6714 | Nup188-Halo Gar1-GFP Pdr5 delta | This study |
| 6782 | Halo-NLS ,Gar-GFP, Pdr5 delta | This study. Halo-NLS, PDR5 delta generated by R.Reyes lab. |
| 6754 | ATTIR1-9myc, MLP1-GFP, Nup188-tdTomato, Rpb2-AID-HA | This study, URA::ADH1-ATTIR1-9myc background generated in Morawska et al, 2013 |
| 6757 | ATTIR1-9myc, MLP1-GFP , Nup188-tdTomato, Rpa135-AID-HA | This study, URA::ADH1-ATTIR1-9myc background generated in Morawska et al, 2013 |
| 6758 | Mex67-5, Nup188-tdTomato Mlp1-GFP | This study |
| 6755 | Rpb1-1, Nup188-tdTomato Mlp1-GFP | This study |
| 6426 | Nup188-tdTomato, Mlp1-GFP | This study |
| 6699 | ATTIR1-9myc, Mlp1GFP, Gar1-tdTomato, Enp1-AID-HA | This study, URA::ADH1-ATTIR1-9myc background generated in Morawska et al, 2013 |
| 6700 | ATTIR1-9myc, Mlp1GFP, Gar1-tdTomato, Csl4-AID-HA | This study, URA::ADH1-ATTIR1-9myc background generated in Morawska et al, 2013 |
| 6720 | ATTIR1-9myc, Mlp1GFP, Gar1-tdTomato, RNA14-AID-HA | This study, URA::ADH1-ATTIR1-9myc background generated in Morawska et al, 2013 |
| 6729 | ATTIR1-9myc, Mlp1GFP, Gar1-tdTomato, RNA15-AID-HA | This study, URA::ADH1-ATTIR1-9myc background generated in Morawska et al, 2013 |
| 6783 | ATTIR1-9myc, Mlp1-prA, Gar1-tdTomato, Enp1-AID-HA | This study, URA::ADH1-ATTIR1-9myc background generated in Morawska et al, 2013 |
| 6721 | ATTIR1-9myc, Mlp1-GFP, Nup60-AID-HA, Gar1-tdTomato | This study, URA::ADH1-ATTIR1-9myc background generated in Morawska et al, 2013 |
| 6748 | ATTIR1-9myc, Mlp1-GFP, Esc1-AID-HA, Gar1-tdTomato | This study, URA::ADH1-ATTIR1-9myc background generated in Morawska et al, 2013 |
| 6766 | ATTIR1-9myc, Mlp1-GFP, Ulp1-AID-HA, Gar1-tdTomato | This study, URA::ADH1-ATTIR1-9myc background generated in Morawska et al, 2013 |
| 6765 | ATTIR1-9myc, Mlp1-GFP, Mlp2-AID-HA | This study, URA::ADH1-ATTIR1-9myc background generated in Morawska et al, 2013 |
| 6731 | ATTIR1-9myc, Mlp1-GFP, Yra1-AID-HA | This study, URA::ADH1-ATTIR1-9myc background generated in Morawska et al, 2013 |
| 6737 | ATTIR1-9myc, Mlp1-GFP, Tho2-AID-HA | This study, URA::ADH1-ATTIR1-9myc background generated in Morawska et al, 2013 |
| 6740 | ATTIR1-9myc, Mlp1-GFP, Sac3-AID-HA | This study, URA::ADH1-ATTIR1-9myc background generated in Morawska et al, 2013 |
| 6742 | ATTIR1-9myc, Mlp1-GFP, Sus1-AID-HA | This study, URA::ADH1-ATTIR1-9myc background generated in Morawska et al, 2013 |
| 6727 | ATTIR1-9myc, Mlp1-GFP, Nab2-AID-HA | This study, URA::ADH1-ATTIR1-9myc background generated in Morawska et al, 2013 |
| 6701 | ATTIR1-9myc, Mlp1-GFP, Pab1-AID-HA, Gar1-tdTomato | This study, URA::ADH1-ATTIR1-9myc background generated in Morawska et al, 2013 |
| 6726 | ATTIR1-9myc, Mlp1-GFP, Pap1-AID-HA | This study, URA::ADH1-ATTIR1-9myc background generated in Morawska et al, 2013 |
| 6722 | ATTIR1-9myc, Mlp1-GFP, Pml39-AID-HA, Gar1-tdTomato | This study, URA::ADH1-ATTIR1-9myc background generated in Morawska et al, 2013 |
| 6724 | ATTIR1-9myc, Mlp1-GFP, Pml1-AID-HA | This study, URA::ADH1-ATTIR1-9myc background generated in Morawska et al, 2013 |
| 6747 | ATTIR1-9myc, Mlp1-GFP, Prp5-AID-HA, Gar1-tdTomato | This study, URA::ADH1-ATTIR1-9myc background generated in Morawska et al, 2013 |
| 6752 | ATTIR1-9myc, Mlp1-GFP, Snu17-AID-HA | This study, URA::ADH1-ATTIR1-9myc background generated in Morawska et al, 2013 |
| 6753 | ATTIR1-9myc, Mlp1-GFP, Luc7-AID-HA | This study, URA::ADH1-ATTIR1-9myc background generated in Morawska et al, 2013 |
| 6784 | ATTIR1-9myc, Mlp1-GFP, Mex67-AID-HA | This study, URA::ADH1-ATTIR1-9myc background generated in Morawska et al, 2013 |
| 6735 | ATTIR1-9myc, Mlp1-GFP, Hrb1-AID-HA | This study, URA::ADH1-ATTIR1-9myc background generated in Morawska et al, 2013 |
| 6728 | ATTIR1-9myc, Mlp1-GFP, Cbc2-AID-HA | This study, URA::ADH1-ATTIR1-9myc background generated in Morawska et al, 2013 |
| 6741 | ATTIR1-9myc, Mlp1-GFP, Gbp2-AID-HA | This study, URA::ADH1-ATTIR1-9myc background generated in Morawska et al, 2013 |
| 6738 | ATTIR1-9myc, Mlp1-GFP,Npl3-AID-HA | This study, URA::ADH1-ATTIR1-9myc background generated in Morawska et al, 2013 |
| 6732 | ATTIR1-9myc, Mlp1-GFP,Prp18-AID-HA | This study, URA::ADH1-ATTIR1-9myc background generated in Morawska et al, 2013 |
| 6746 | ATTIR1-9myc, MLP1-GFP, Gar1-tdTomato, Rpb2-AID-HA | This study, URA::ADH1-ATTIR1-9myc background generated in Morawska et al, 2013 |
| 6683 | Nup188-tdTomato, Nup84-GFP | This study |
| 6785 | Nup188-GFP, Gar1-tdTomato | This study |
| 6881 | Nup188-tdTomato, Nup133-GFP | This study |
| 6682 | Nup188-tdTomato, Nup49-GFP | This study |
| 6687 | Nup188-tdTomato, Nup60-GFP | This study |
| 6685 | Nup188-tdTomato, Sac3-GFP | This study |
| 6688 | Nup188-tdTomato, Pml39-GFP | This study |
| 6686 | Nup188-tdTomato, Ulp1-GFP | This study |
| 6689 | Nup188-tdTomato, Mex67-GFP | This study |
| 6786 | Mlp1-Halo, Mlp2-GFP | This study |
| 6696 | Mlp1-Halo, Sac3-GFP | This study |
| 6698 | Mlp1-Halo, Pml39-GFP | This study |
| 6697 | Mlp1-Halo, Mex67-GFP | This study |
| 6643 | Mlp1-Halo, Ulp1-GFP | This study |
| 6717 | Mlp1-GFP, Sac3-Halo | This study |
| 6787 | Mlp1-GFP, Pml39-Halo | This study |
| 6716 | Mlp1-GFP, Ulp1-Halo | This study |
| 6769 | ATTIR1-9myc, MLP1-GFP, Rpb2-AID-HA, Pml39-tdTomato | This study |
| 6768 | ATTIR1-9myc, MLP1-GFP, Rpb2-AID-HA, Ulp1-tdTomato | This study |
| 6688 | DeltaMlp1/Mlp2 Nup133 GFP | This study |
