## Supplementary material for "Nuclear mRNA metabolism drives selective basket assembly on a subset of nuclear pores in budding yeast": Table S3

**Table S3. Primers for construct generation**

| Name | Sequence |
|---|---|
| Mlp1-GFP (Fwd) | AAGATGAGGAAGAAAAAGAAACCGATAAGGTGAATGACGAGAACAGTATACGTACGCTGCAGGTCGAC |
| Mlp1-GFP (Rev) | ACATTGAAAAAGGTTTAGTTTGTATTGATCCCTTGTTTTTACTATCTCCTATCGATGAATTCGAGCTCG |
| Gar1-tdTomato (Fwd) | GGATCTCGTGGCGGATCTCGTGGTGGTTTCAGAGGAGGTAGAAGAGCTCTAGAACTAGTGGATCC |
| Gar1-tdTomato (Rev) | CAGATATAGTAAGTTGGAAGAAATGAAGAATTGTGAAAGATAAAGGGCATAGGCCACTAGTGGATCTG |
| Nup188-tdTomato (Fwd) | CAAGGGTATCAGCAGAGACATTAAAGCATTACAAGATTCACTATTTAAGGACGTTGCTCTAGAACTAGTGGATCC |
| Nup188-tdTomato (Rev) | GCACTTGCACTGTTCATTATTATATTATGTAGCTTTACATAACCTGCAAAATAAGGCATAGGCCACTAGTGGATCTG |
| Mlp1-Halo (Fwd) | GAGGAAGAAAAAGAAACCGATAAGGTGAATGACGAGAACAGTATAGGTGACGGTGCTGGTTTA |
| Mlp1-Halo (Rev) | GCAGAATGAAGCTCCTCCACATTGAAAAAGGTTTAGTTTGTATTGACACAGGAAACAGCTATGACC |
| Nup188-Halo (Fwd) | CAAGGGTATCAGCAGAGACATTAAAGCATTACAAGATTCACTATTTAAGGACGTTGGTGACGGTGCTGGTTTA |
| Nup188-Halo (Rev) | GCACTTGCACTGTTCATTATTATATTATGTAGCTTTACATAACCTGCAAAATAAGACACAGGAAACAGCTATGACC |
| Rpa135-AID-HA (Fwd) | CTATCCGCAATGGGTATAAGATTGCGTTATAATGTAGAGCCCAAACGTACGCTGCAGGTCGAC |
| Rpa135-AID-HA (Rev) | CCTTCATTTACCATTCTATATCAATTTGGAAAGAAGGGTATTTCTATCGATGAATTCGAGCTCG |
| Rpb2-AID-HA (Fwd) | ATGAACATTACACCACGTTTATATACCGATCGTTCGAGAGATTTTCGTACGCTGCAGGTCGAC |
| Rpb2-AID-HA (Rev) | AATGTTTTTTATTATTTTACTTTCTTAGAGTTACAACATTATTTCATCGATGAATTCGAGCTCG |
| Enp1-AID-HA (Fwd) | CAGGGAGTTTGTTGATCCACAGGAAGCTAATGATGATTTAATGATTGATGTCAATCGTACGCTGCAGGTCGAC |
| Enp1-AID-HA (Rev) | TGAAAGGGGGAAAGACCGAGCGATATAAAATTGATGAAAAATTGATATTACAGCAATCGATGAATTCGAGCTCG |
| Csl4-AID-HA (Fwd) | GATGACTTCACCGGTTACAGGCGCTACAGAAAAGCGCAAATGTGCCAAACCTTTTCGTACGCTGCAGGTCGAC |
| Csl4-AID-HA (Rev) | TACCCTCTTTTAAATATATACGCGTCTATATGCACTGTAGATAAGCTGTTACATAATCGATGAATTCGAGCTCG |
| Rna14-AID-HA (Fwd) | GAATTTTTTAAATGATCAAGTAGAGATTCCAACAGTTGAGAGCACCAAGTCAGGTCGTACGCTGCAGGTCGAC |
| Rna14-AID-HA (Rev) | TTATAATAGATGTGTTGGTATAAATATTCATATATACCTATTTATTAACGTAATGATCGATGAATTCGAGCTCG |
| Rna15-AID-HA (Fwd) | GATGGCTATTTGGGACTTAAAACAAAAAGCATTAAGGGGAGAATTTGGTGCATTTCGTACGCTGCAGGTCGAC |
| Rna15-AID-HA (Rev) | GTTGCCTCATCATTGCGGAACCGCATTTTTTTTTTGTATTTTTGCCTCCCTAGTTATCGATGAATTCGAGCTCG |
| Nup60-AID-HA (Fwd) | AAATGGCTTGGTTGATGAAAATAAAGTTGAGGCTTTCAAGTCCCTATATACCTTTCGTACGCTGCAGGTCGAC |
| Nup60-AID-HA (Rev) | CTTACGTATTGAGTTGGGCTATACGGTAATTATGTCACGGCTAAAATTTTCATTAATCGATGAATTCGAGCTCG |
| Esc1-AID-HA (Fwd) | TAGGGGGCACGAGCCAAAAAGCCGTGGACAGAATACGCATCCAAGTGTTGACAAACGTACGCTGCAGGTCGAC |
| Esc1-AID-HA (Rev) | AGAAAAACGCATCGCAATAATTATTACTATCTACATATTCCTGTATACAATTTGAATCGATGAATTCGAGCTCG |
| Ulp1-AID-HA (Fwd) | TGCGATTAGGATGAGAAGATTTATTGCCCATTTGATTTTAACCGACGCTTTAAAACGTACGCTGCAGGTCGAC |
| Ulp1-AID-HA (Rev) | CAATGATCTGAATATTTCTACTTATGTATAATAATTGTATATTATAAAAGAATAAATCGATGAATTCGAGCTCG |
| Mlp2-AID-HA (Fwd) | ACACCAAAAAGGTTAAAGAGAGTCCAGCAAATGATCAAGCTTCCAACGAGCGTACGCTGCAGGTCGAC |
| Mlp2-AID-HA (Rev) | AAAATATGTAGATGTTTCATATTTATATAATTACATTGTTTAATATTACAATCGATGAATTCGAGCTCG |
| Yra1-AID-HA (Fwd) | TAAGAAAAGTCTTGAAGATCTGGACAAGGAAATGGCGGACTATTTCGAAAAGAAACGTACGCTGCAGGTCGAC |
| Yra1-AID-HA (Rev) | GGaaaaattaaatttaataaaaccaaattaaatcaaacaaaaaaTTGACAATTAAATCGATGAATTCGAGCTCG |
| Tho2-AID-HA (Fwd) | TCAGGCGCTTCCGCAAGGTCCCAAGGGTGGGAATTACGTCAGTAGGTACCAGAGGCGTACGCTGCAGGTCGAC |
| Tho2-AID-HA (Rev) | GGGAACTATCAAAGTACACGTTAAAATTCAGCTCGGGTATGTTAAGTACTAGTAAATCGATGAATTCGAGCTCG |
| Sac3-AID-HA (Fwd) | TATATTAGAGCTGAAGATCTTGATCGATTCTGTCAAGAAGAAAGTAAATAATGATCGTACGCTGCAGGTCGAC |
| Sac3-AID-HA (Rev) | TTCCTAAAGCTATAGAAAAAATGCACATTTCTTTTGTTTATATATTACAAATGCTATCGATGAATTCGAGCTCG |
| Sus1-AID-HA (Fwd) | GTTTTAAAGCAAATAAGGGAATTTCTTGAAGAGATTGTAGATACACAACGTACGCTGCAGGTCGAC |
| Sus1-AID-HA (Rev) | TTTCCCGATGAGCATATGTAATAATATTGGGAATTAAGGTGCATTTTCGTATCCTATCGATGAATTCGAGCTCG |
| Nab2-AID-HA (Fwd) | AAATGCTCCTCCGCAAACCAGTTTTACGCACCAAGAACAAGATACGGAAATGAACCGTACGCTGCAGGTCGAC |
| Nab2-AID-HA (Rev) | CTTCCATCAAAAGGGTCACAGGAACATGAATTTCGTTCCGTGATTTTAATAGTAAATCGATGAATTCGAGCTCG |
| Pab1-AID-HA (Fwd) | TTCTGCTGCCTATGAGTCTTTCAAAAAGGAGCAAGAACAACAAACTGAGCAAGCTCGTACGCTGCAGGTCGAC |
| Pab1-AID-HA (Rev) | AGAAAAAAAAGATGATAAGTTTGTTGAGTAGGGAAGTAGGTGATTACATAGAGCAATCGATGAATTCGAGCTCG |
| Pap1-AID-HA (Fwd) | AGATGCTGCTTCAGGTGACAACATCAATGGCACAACCGCAGCTGTTGACGTAAACCGTACGCTGCAGGTCGAC |
| Pap1-AID-HA (Rev) | GTTTATGACTGATTAACCTATATTAATAAACTATTCAACTATAAATAGGAATGTCATCGATGAATTCGAGCTCG |
| Pml39-AID-HA (Fwd) | GAAATTGGGCGTGGGAGAAAGACTAAATAAATTAGAGGCTGTTCTACAAACTTTACGTACGCTGCAGGTCGAC |
| Pml39-AID-HA (Rev) | CAGCATGGGGGCATATACAAGCATATGAGAATTTGGATAATGTATTACATCTAATATCGATGAATTCGAGCTCG |
| Pml1-AID-HA (Fwd) | TACACTTTCAGAATTTGAAGAAGATACCGATTACGAACTCATCTTCATGAATGTACGTACGCTGCAGGTCGAC |
| Pml1-AID-HA (Rev) | CAGCATTCAAAGAAGAATAATTAAAACACACTGAAAGTGTGTTTCTTATATATGGATCGATGAATTCGAGCTCG |
| Prp5-AID-HA (Fwd) | GGGGTCGTAAAGGCTGCAAGCTTGTCTTTGAAGAGTACTAAATACCGTACGCTGCAGGTCGAC |
| Prp5-AID-HA (Rev) | AACTACGAAAGTATATAGCACCACGAGTGAGTTAAATTCTAAAAAATCGATGAATTCGAGCTCG |
| Snu17-AID-HA (Fwd) | ATAGCTGATAGACTGTGGAGTCGTAAAGAATTTCGCTTGGGGACCCGTACGCTGCAGGTCGAC |
| Snu17-AID-HA (Rev) | GAGCGAGCCTTTCCCTTTTGGGACGCGCGCCAAGGCCCTTCTGTTATCGATGAATTCGAGCTCG |
| Luc7-AID-HA (Fwd) | AACGCCAGCAAGACAGCTACTACACTACCCGGAAGACGCTTTGTGCGTACGCTGCAGGTCGAC |
| Luc7-AID-HA (Rev) | TCCTTCGAACAAAATTTTTCTAGCATCATTTTTTTATGTATGGCCATCGATGAATTCGAGCTCG |
| Mex67-AID-HA (Fwd) | AAAGGGTTTTCAGAGTAGCATGAATGGCATCCCTAGAGAAGCATTTGTGCAGTTCCGTACGCTGCAGGTCGAC |
| Mex67-AID-HA (Rev) | GCTTAAACTGTATATTTTTTGTGATACTGTGCGGCTGAAACAGGGAACAATATCAATCGATGAATTCGAGCTCG |
| Hrb1-AID-HA (Fwd) | GAATAATTATAACTATGGGGGTTGTGATTTGGATATATCGTACGCTAAACGCCTCCGTACGCTGCAGGTCGAC |
| Hrb1-AID-HA (Rev) | ATAAATACTTGTCGCAGATCCAATAGGTGAGAAAGTATATAGATCGAGAGTAGTTATCGATGAATTCGAGCTCG |
| Cbc2-AID-HA (Fwd) | TACTTTCAGACCAGGTTTCGATGAAGAAAGAGAAGATGATAACTACGTACCTCAGCGTACGCTGCAGGTCGAC |
| Cbc2-AID-HA (Rev) | atatatatatatatatCTGTGTGTAGAATCTTTCTCAGATATAAATTGATTGATTATCGATGAATTCGAGCTCG |
| Gbp2-AID-HA (Fwd) | AAATAATTATAATTATGGTGGTTGTAGTTTACAGATCTCTTATGCTAGACGTGATCGTACGCTGCAGGTCGAC |
| Gbp2-AID-HA (Rev) | TTATTTATACGTTATCATAAAGTACACAGGTCATGGTTCGGTTGGTGCTTAGGAAATCGATGAATTCGAGCTCG |
| Npl3-AID-HA (Fwd) | TCCAAGAGATGCATACAGAACCAGAGATGCTCCACGTGAAAGATCACCAACCAGGCGTACGCTGCAGGTCGAC |
| Npl3-AID-HA (Rev) | ACAATTCATATCTTTTGTTAATTTCTCCTTTTTTTTTCTCAACTATATAAATGGCATCGATGAATTCGAGCTCG |
| Prp18-AID-HA (Fwd) | TAAAAGATTAATAACTTTTGAAGAATGGTATACCAGCAACCACGATAGCTTAGCCCGTACGCTGCAGGTCGAC |
| Prp18-AID-HA (Rev) | TTATTTTGGCCGCATGATATCGTGCCACGCGATAACGAAAACAATAGTTCAACAAATCGATGAATTCGAGCTCG |
| Nup84-GFP (Fwd) | TGGAAAGTTAAAAGAGTATCTGGATCTCGTTGCTCGCACAGCAACCCTTTCGAACCGTACGCTGCAGGTCGAC |
| Nup84-GFP (Rev) | TAAAATTATTGCTGTTTACTTAAAATATAAACTTATTCTGCAATACATTAATTGAATCGATGAATTCGAGCTCG |
| Nup60-GFP (Fwd) | AAATGGCTTGGTTGATGAAAATAAAGTTGAGGCTTTCAAGTCCCTATATACCTTTCGTACGCTGCAGGTCGAC |
| Nup60-GFP (Rev) | CTTACGTATTGAGTTGGGCTATACGGTAATTATGTCACGGCTAAAATTTTCATTAATCGATGAATTCGAGCTCG |
| Nup49-GFP (Fwd) | GCCGTGTTACATCAAAAAACGAAAACACTGGCATCATTGAGCATAGCTCTAGAACTAGTGGATCC |
| Nup49-GFP (Rev) | TGTACAAGACATTTGTACTTGTTATACGCACTATATAAACTTTCAGCATAGGCCACTAGTGGATCTG |
| Nup133-GFP (Fwd) | TGTAGCGAAAGAAAAAAACTATACCATCAACTATGAAACCAACACTGTAGAATACCGTACGCTGCAGGTCGAC |
| Nup133-GFP (Rev) | TATTATCATTCCCCAGTAAAGTTTATTATATATATGTAAAATTGTATTATAGATAATCGATGAATTCGAGCTCG |
| Ulp1-GFP (Fwd) | TGCGATTAGGATGAGAAGATTTATTGCCCATTTGATTTTAACCGACGCTTTAAAACGTACGCTGCAGGTCGAC |
| Ulp1-GFP (Rev) | CAATGATCTGAATATTTCTACTTATGTATAATAATTGTATATTATAAAAGAATAAATCGATGAATTCGAGCTCG |
| Pml39-GFP (Fwd) | AGGAGAAATAAAACATTATTCCCCAGGAATTGAGAGGAAAGTAGGGCAGTTACTACGTACGCTGCAGGTCGAC |
| Pml39-GFP (Rev) | CAGCATGGGGGCATATACAAGCATATGAGAATTTGGATAATGTATTACATCTAATATCGATGAATTCGAGCTCG |
| Mex67-GFP (Fwd) | AAAGGGTTTTCAGAGTAGCATGAATGGCATCCCTAGAGAAGCATTTGTGCAGTTCCGTACGCTGCAGGTCGAC |
| Mex67-GFP (Rev) | GCTTAAACTGTATATTTTTTGTGATACTGTGCGGCTGAAACAGGGAACAATATCAATCGATGAATTCGAGCTCG |
| Sac3-GFP (Fwd) | TATATTAGAGCTGAAGATCTTGATCGATTCTGTCAAGAAGAAAGTAAATAATGATCGTACGCTGCAGGTCGAC |
| Sac3-GFP (Rev) | TTCCTAAAGCTATAGAAAAAATGCACATTTCTTTTGTTTATATATTACAAATGCTATCGATGAATTCGAGCTCG |
| Mlp2-GFP (Fwd) | ACACCAAAAAGGTTAAAGAGAGTCCAGCAAATGATCAAGCTTCCAACGAGCGTACGCTGCAGGTCGAC |
| Mlp2-GFP (Rev) | AAAATATGTAGATGTTTCATATTTATATAATTACATTGTTTAATATTACAATCGATGAATTCGAGCTCG |
| Ulp1-Halo (Fwd) | TGCGATTAGGATGAGAAGATTTATTGCCCATTTGATTTTAACCGACGCTTTAAAAGGTGACGGTGCTGGTTTA |
| Ulp1-Halo (Rev) | CAATGATCTGAATATTTCTACTTATGTATAATAATTGTATATTATAAAAGAATAAACACAGGAAACAGCTATGACC |
| Pml39-Halo (Fwd) | GAAATTGGGCGTGGGAGAAAGACTAAATAAATTAGAGGCTGTTCTACAAACTTTAGGTGACGGTGCTGGTTTA |
| Pml39 Halo (Rev) | CAGCATGGGGGCATATACAAGCATATGAGAATTTGGATAATGTATTACATCTAATACACAGGAAACAGCTATGACC |
| Sac3-Halo (Fwd) | TATATTAGAGCTGAAGATCTTGATCGATTCTGTCAAGAAGAAAGTAAATAATGATGGTGACGGTGCTGGTTTA |
| Sac3-Halo (Rev) | TTCCTAAAGCTATAGAAAAAATGCACATTTCTTTTGTTTATATATTACAAATGCTACACAGGAAACAGCTATGACC |
| Nab2-tdTomato (Fwd) | AAATGCTCCTCCGCAAACCAGTTTTACGCACCAAGAACAAGATACGGAAATGAACGCTCTAGAACTAGTGGATCC |
| Nab2-tdTomato (Rev) | TTGAATAGGTGTCTTCCATCAAAAGGGTCACAGGAACATGAATTTCGTTCCGTGAGCATAGGCCACTAGTGGATCTG |
| Sac3-tdTomato (Fwd) | TATATTAGAGCTGAAGATCTTGATCGATTCTGTCAAGAAGAAAGTAAATAATGATGCTCTAGAACTAGTGGATCC |
| Sac3-tdTomato (Rev) | TTCCTAAAGCTATAGAAAAAATGCACATTTCTTTTGTTTATATATTACAAATGCTGCATAGGCCACTAGTGGATCTG |
| Pml39-tdTomato (Fwd) | GAAATTGGGCGTGGGAGAAAGACTAAATAAATTAGAGGCTGTTCTACAAACTTTAGCTCTAGAACTAGTGGATCC |
| Pml39-tdTomato (Rev) | CAGCATGGGGGCATATACAAGCATATGAGAATTTGGATAATGTATTACATCTAATGCATAGGCCACTAGTGGATCTG |
| Ulp1-tdTomato (Fwd) | TGCGATTAGGATGAGAAGATTTATTGCCCATTTGATTTTAACCGACGCTTTAAAAGCTCTAGAACTAGTGGATCC |
| Ulp1-tdTomato (Rev) | CAATGATCTGAATATTTCTACTTATGTATAATAATTGTATATTATAAAAGAATAAGCATAGGCCACTAGTGGATCTG |
| Mlp1-PrA(Fwd) | GACTGAAGATGAGGAAGAAAAAGAAACCGATAAGGTGAATGACGAGAACAGTATAGGTGAAGCTCAAAAACTTAAT |
| Mlp1-PrA(Rev) | CCTCCACATTGAAAAAGGTTTAGTTTGTATTGATCCCTTGTTTTTACTATCTCCTATCGATGAATTCGAGCTCG |
