## Supplementary material for "Nuclear mRNA metabolism drives selective basket assembly on a subset of nuclear pores in budding yeast": Table S4

**Table S4. Plasmids**

| Description | Reference |
|---|---|
| centromeric plasmid pEXPGFPNLS_M2NT1 for Nterm1 Mlp1 fragment expression, HIS3 | Niepel et al., 2013 |
| centromeric plasmid pEXPGFPNLS_M2NT2 for Nterm2 Mlp1 fragment expression, HIS3 | Niepel et al., 2013 |
| centromeric plasmid pEXPGFPNLS_M2NT3 for Nterm3 Mlp1 fragment expression, HIS3 | Niepel et al., 2013 |
| centromeric plasmid pEXPGFPNLS_M2NT4 for Nterm4 Mlp1 fragment expression, HIS3 | Niepel et al., 2013 |
| centromeric plasmid pEXPGFPNLS_M2NT5 for Nterm5 Mlp1 fragment expression, HIS3 | Niepel et al., 2013 |
| centromeric plasmid pEXPGFPNLS_M2NT6 for Nterm6 Mlp1 fragment expression, HIS3 | Niepel et al., 2013 |
| centromeric plasmid pEXPGFPNLS_M2CT for Cterm Mlp1 fragment expression, HIS3 | Niepel et al., 2013 |
| centromeric plasmid pUG34_GFPNLS HIS3 | Niepel et al., 2013 |
| pZUT3 centromeric plasmid carrying GAR1-GFP, URA3 | Trumtel et al., 2000 |
