## Supplementary material for "Nuclear mRNA metabolism drives selective basket assembly on a subset of nuclear pores in budding yeast": Table S6

**Table S6. Processed AP-MS data**

| Alternate ID | Basket-Plus (Mlp1 PrA) | Basket-minus (Nup133-GFP) | All NPCs (Nup133-GFP) | Delta *mlp1/2* NPCs (Nup133-GFP) | Enp1-AID-HA +auxin (Mlp1-PrA) | Complex |
|---|---|---|---|---|---|---|
| **MLP1** | 2017.001828 | 9.715994021 | 135.7923497 | 0 | 578.7037037 | Basket |
| **NSP1** | 145.0335161 | 38.56502242 | 33.60655738 | 20.74592075 | 63.88888889 | Central core |
| **NUP192** | 112.4314442 | 29.59641256 | 44.80874317 | 21.21212121 | 103.7037037 | Central core |
| **NUP96** | 107.8610603 | 31.39013453 | 37.15846995 | 16.78321678 | 73.14814815 | Central core |
| **NUP145** | 104.0219378 | 65.76980568 | 80.87431694 | 62.7039627 | 84.25925926 | Central core |
| **MLP2** | 100 | 0.597907324 | 18.30601093 | 0 | 100.9259259 | Basket |
| **Pom152** | 99.08592322 | 28.25112108 | 36.33879781 | 20.51282051 | 86.11111111 | Central core |
| **NSP1** | 92.13893967 | 26.00896861 | 16.12021858 | 7.692307692 | 23.14814815 | Central core |
| **NUP157** | 80.31687995 | 24.96263079 | 33.06010929 | 17.71561772 | 76.85185185 | Central core |
| **Nup133** | 72.02925046 | 100.1494768 | 100.273224 | 100 | 37.96296296 | Central core |
| **NUP188** | 68.73857404 | 24.96263079 | 36.8852459 | 18.18181818 | 77.77777778 | Central core |
| **Nup159** | 68.73857404 | 19.43198804 | 22.13114754 | 9.324009324 | 29.62962963 | Central core |
| **NUP85** | 65.26508227 | 38.71449925 | 32.24043716 | 22.61072261 | 12.03703704 | Central core |
| **NUP120** | 58.86654479 | 47.08520179 | 37.43169399 | 27.73892774 | 32.40740741 | Central core |
| **NUP120** | 54.66179159 | 42.3019432 | 32.24043716 | 24.94172494 | 32.40740741 | Central core |
| **NUP2** | 53.3820841 | 2.541106129 | 7.650273224 | 0 | 31.48148148 | Basket |
| **NUp82** | 46.6179159 | 14.34977578 | 17.21311475 | 10.72261072 | 22.22222222 | Central core |
| **NUP84** | 44.60694698 | 65.76980568 | 68.85245902 | 59.67365967 | 29.62962963 | Central core |
| **NUP60** | 42.59597806 | 1 | 6.557377049 | 0 | 41.66666667 | Basket |
| **NUP57** | 36.92870201 | 14.64872945 | 18.85245902 | 10.02331002 | 33.33333333 | Central core |
| **NUP116** | 34.73491773 | 13.30343797 | 15.0273224 | 12.12121212 | 30.55555556 | Central core |
| **NUP100** | 32.54113346 | 7.623318386 | 9.836065574 | 4.895104895 | 19.44444444 | Central core |
| **ASM4** | 29.61608775 | 6.726457399 | 8.196721311 | 1.631701632 | 9.259259259 | Central core |
| **SEC13** | 21.81596587 | 21.97309417 | 25.13661202 | 21.67832168 | 43.51851852 | Central core |
| **GLE2** | 18.09872029 | 7.174887892 | 4.371584699 | 1.864801865 | 5.555555556 | Central core |
| **NUP85** | 17.55027422 | 6.128550075 | 4.644808743 | 3.263403263 | 1.851851852 | Central core |
| **POM34** | 15.72212066 | 2.242152466 | 1.912568306 | 0.932400932 | 5.555555556 | Central core |
| **NDC1** | 15.35648995 | 7.772795217 | 6.284153005 | 3.03030303 | 15.74074074 | Central core |
| **NUP157** | 13.89396709 | 4.783258595 | 3.551912568 | 2.564102564 | 12.96296296 | Central core |
| **NUP145** | 12.06581353 | 7.025411061 | 10.92896175 | 6.75990676 | 7.407407407 | Central core |
| **NUP1** | 4.387568556 | 0.4 | 2.5 | 0 | 0 | Basket |
| **NUP84** | 3.65630713 | 4.334828102 | 5.464480874 | 3.962703963 | 2.777777778 | Central core |
| **GLE1** | 2.559414991 | 0.597907324 | 1.639344262 | 0.932400932 | 5.555555556 | Central core |
| **Nup42** | 0.914076782 | 0 | 0 | 0 | 0 | Central core |
| **Nup53** | 42.23034735 | 12.85500747 | 15.0273224 | 5.594405594 | 25 | Karyopherins & importin |
| | 1718.464351 | 728.7542601 | 930.9153005 | 519.8135198 | 1756.481481 | |
| | 36 | 35 | 35 | 29 | 34 | |
| **Kap95** | 43.20536258 | 4.334828102 | 9.836065574 | 3.962703963 | 50.92592593 | Karyopherins & importin |
| **SRP1** | 25.65508836 | 0 | 2.459016393 | 0 | 12.03703704 | Karyopherins & importin |
| **Kap123** | 11.2126752 | 2.840059791 | 7.37704918 | 4.428904429 | 25 | Karyopherins & importin |
| **KAP120** | 6.398537477 | 0 | 0.273224044 | 0 | 0 | Karyopherins & importin |
| **Pse1** | 0.914076782 | 0.149476831 | 0.546448087 | 3.03030303 | 12.96296296 | Karyopherins & importin |
| **MTR10** | 0.365630713 | 0 | 0 | 0 | 0 | Karyopherins & importin |
| **RNA1** | 0.060938452 | 1.644245142 | 4.918032787 | 3.962703963 | 25.92592593 | Karyopherins & importin |
| | 87.81230957 | 8.968609865 | 25.40983607 | 15.38461538 | 126.8518519 | |
| | 7 | 4 | 6 | 5 | 6 | |
| **MEX67** | 182.2669104 | 20.62780269 | 23.7704918 | 12.58741259 | 55.55555556 | export receptor |
| **YRA1** | 17.97684339 | 3.736920777 | 3.819672131 | 1.165501166 | 41.66666667 | Trex |
| **PAB1** | 14.07678245 | 10.91180867 | 5.737704918 | 7.692307692 | 101.8518519 | PolyA binding protein |
| **PML39** | 12.6142596 | 0 | 1.639344262 | 0 | 4.62962963 | Pml39 |
| **STO1** | 9.323583181 | 0 | 0 | 0 | 6.481481481 | Cap |
| **YHR127W** | 6.032906764 | 1.34529148 | 1.092896175 | 0 | 10.18518519 | ND |
| **MTR2** | 4.753199269 | 2.242152466 | 4.371584699 | 1.398601399 | 12.03703704 | export receptor |
| **NAB2** | 3.10786106 | 0 | 0.273224044 | 0 | 1.851851852 | PolyA binding protein |
| **Yra2** | 2.437538087 | 0.448430493 | 1.092896175 | 0.932400932 | 2.777777778 | polyA export |
| **Hpr1** | 0.731261426 | 0.149476831 | 1.092896175 | 0.466200466 | 0.925925926 | THO complex |
| **TIF4632** | 0.670322974 | 0.747384155 | 0 | 0 | 0 | cap |
| **Los1** | 0.670322974 | 0 | 1.092896175 | 0.466200466 | 0.925925926 | cap |
| **CBC2** | 0.365630713 | 0.448430493 | 0 | 0 | 0 | cap |
| **Tho2** | 0.365630713 | 0 | 0.819672131 | 1.165501166 | 7.407407407 | THO complex |
| **Gbp2** | 0 | 0.597907324 | 1.639344262 | 4.428904429 | 18.51851852 | HnRnp |
| **CDC31** | 17.00182815 | 0.2 | 3.825136612 | 2.797202797 | 13.88888889 | Trex2 |
| **FIR1** | 0.182815356 | 0.448430493 | 4.918032787 | 1.631701632 | 0.925925926 | 3'RNA processing |
| | 272.5776965 | 41.90403587 | 55.18579235 | 34.73193473 | 279.6296296 | |
| | 20 | 15 | 18 | 12 | 20 | |
| **SAC3** | 68.43388178 | 1.195814649 | 13.38797814 | 2.097902098 | 63.88888889 | Trex2 |
| **SUS1** | 10.42047532 | 0.149476831 | 3.551912568 | 0 | 12.96296296 | Trex2 |
| **THP1** | 9.87202925 | 0.149476831 | 1.912568306 | 0 | 7.407407407 | Trex2 |
| **Sub2** | 4.936014625 | 0 | 4.918032787 | 0 | 20.37037037 | Trex2 |
| **CDC31** | 17.00182815 | 0.2 | 3.825136612 | 2.797202797 | 13.88888889 | Trex2 |
| **PML39** | 12.6142596 | 0 | 1.639344262 | 0 | 4.62962963 | Pml39 |
| **ULP1** | 2.742230347 | 0.149476831 | 0.546448087 | 0 | 1.851851852 | Central core |
| **MAD1** | 8.592321755 | 0 | 0.7 | 0 | 12.96296296 | SPD |
| **MAD2** | 2.010968921 | 0 | 0 | 0 | 0 | SPD |
| **MPS3** | 1.462522852 | 0 | 0.819672131 | 0 | 0 | SPD |
| **MPS2** | 0.731261426 | 0 | 0 | 0 | 0 | SPD |
| **SPC110** | 0.548446069 | 0.149476831 | 5.191256831 | 0.699300699 | 10.74074074 | SPD |
| **KAR3** | 0.365630713 | 0 | 0 | 0 | 0 | SPD |
| **Kre28** | 0.182815356 | 0 | 0 | 0 | 0 | SPD |
| **Stu1** | 0 | 0.448430493 | 0.273224044 | 1.165501166 | 4.62962963 | SPD |
| **SPC42** | 0 | 0 | 0 | 0.233100233 | 4.62962963 | SPD |
| **CDC31** | 17.00182815 | 0.747384155 | 3.825136612 | 2.797202797 | 13.88888889 | SPD |
| **HOS3** | 3.290676417 | 0 | 2.459016393 | 4.195804196 | 25.92592593 | histone modification |
| **RNR2** | 3.168799512 | 0 | 4.098360656 | 15.38461538 | 22.22222222 | DNA damage control |
| **msh2** | 0.731261426 | 0 | 0.3 | 0 | 0 | Telomere associated protein |
| **Irr1p** | 0.731261426 | 0 | 0.273224044 | 0.233100233 | 2.777777778 | Chromatin organization |
| **Mcm16** | 0.731261426 | 0 | 0 | 0 | 0 | DNA replication |
| **Gcr1** | 0.731261426 | 0 | 0 | 0 | 0 | polII |
| **CTF18** | 0.548446069 | 0.896860987 | 2.18579235 | 1.864801865 | 11.11111111 | DNA replication |
| **Sid2** | 0.548446069 | 0.149476831 | 0 | 0 | 0 | DNA replication |
| **HDA3** | 0.365630713 | 0.298953662 | 0 | 0 | 0 | histone modification |
| **SET2** | 0.182815356 | 1.195814649 | 3.551912568 | 3.962703963 | 0 | histone modification |
| **TOM1** | 0.609384522 | 0 | 0 | 0 | 0 | histone modification |
| **MRC1** | 0.609384522 | 0 | 0 | 0 | 0 | DNA replication |
| **SPT6** | 0.365630713 | 0 | 0 | 0.466200466 | 4.62962963 | histone modification |
| **BRE1** | 0.548446069 | 0.298953662 | 1.912568306 | 1.165501166 | 7.407407407 | histone modification |
| | 13.16270567 | 2.840059791 | 14.78087432 | 27.27272727 | 74.07407407 | |
| | 14 | 5 | 7 | 7 | 6 | |
| **RIF1** | 0.182815356 | 0 | 0 | 0 | 0 | Telomere associated protein |

| | | | | | | |
|---|---|---|---|---|---|---|
| **Cdc48** | 0.182815356 | 0.149476831 | 0.273224044 | 0.233100233 | 0 | Telomere associated protein |
| **msh2** | 0.731261426 | 0 | 0.3 | 0 | 0 | Telomere associated protein |
| | 1.096892139 | 0.149476831 | 0.573224044 | 0.233100233 | 0 | |
| | 3 | 2 | 3 | 2 | 0 | |
| **RPB1** | 2 | 0.7 | 1.6 | 5.8 | 3.7 | polII |
| **Swi6** | 0.365630713 | 0.896860987 | 1.366120219 | 0 | 3.703703704 | transcription cofactor |
| **Taf2** | 0.365630713 | 0 | 0 | 0 | 0 | transcription cofactor |
| **RPB2** | 0.121876904 | 2.092675635 | 3.005464481 | 3.263403263 | 4.62962963 | PolII |
| **SUB1** | 0.060938452 | 0 | 1.366120219 | 1.398601399 | 10.18518519 | transcription cofactor |
| **SPO14** | 0 | 2.989536622 | 8.196721311 | 0.699300699 | 3.703703704 | transcription cofactor |
| **Rpb11** | 0.365630713 | 0.597907324 | 1.092896175 | 0 | 0.925925926 | PolII |
| **RPB9** | 0.182815356 | 0 | 0 | 0 | 0 | PolII |
| **WTM1** | 10.78610603 | 0 | 4.918032787 | 4.662004662 | 30.55555556 | transcription cofactor |
| **RPB3** | 0.792199878 | 0.149476831 | 0 | 0 | 0 | polII |
| **Gcr1** | 0.731261426 | 0 | 0 | 0 | 0 | polII |
| | 15.77209019 | 7.426457399 | 21.54535519 | 15.82331002 | 57.4037037 | |
| | 10 | 6 | 7 | 5 | 7 | |
| **SMD1** | 1.279707495 | 0 | 2.18579235 | 1.165501166 | 9.259259259 | spliceosomal complex |
| **prp9** | 0.548446069 | 1.34529148 | 2.459016393 | 0.233100233 | 3.703703704 | spliceosomal complex |
| **prp45** | 0.548446069 | 0 | 0 | 0 | 0 | spliceosomal complex |
| **SME1** | 0.548446069 | 0 | 0 | 0 | 0 | spliceosomal complex |
| **Luc7** | 0.548446069 | 0 | 0 | 0 | 1.851851852 | spliceosomal complex |
| **prp43** | 0.2 | 0 | 0 | 0 | 1.851851852 | spliceosomal complex |
| | 3.673491773 | 1.34529148 | 4.644808743 | 1.398601399 | 16.66666667 | |
| | 6 | 1 | 2 | 2 | 4 | |
| **RPT5** | 6.642291286 | 6.427503737 | 6.010928962 | 4.195804196 | 6.481481481 | proteasome |
| **SEM1** | 5.850091408 | 0.896860987 | 1.639344262 | 0 | 4.62962963 | proteasome |
| **RPT2** | 5.118829982 | 2.989536622 | 2.732240437 | 0.233100233 | 13.88888889 | proteasome |
| **RPT3** | 4.753199269 | 1.046337818 | 1.639344262 | 2.564102564 | 0 | proteasome |
| **RPT6** | 3.778184034 | 5.381165919 | 8.743169399 | 7.925407925 | 27.77777778 | proteasome |
| **Rpn5** | 2.193784278 | 0 | 0.273224044 | 0 | 0.925925926 | proteasome |
| **PRE4** | 2.071907374 | 0 | 0.546448087 | 0 | 5.555555556 | proteasome |
| **RPN12** | 2.010968921 | 1.195814649 | 0.273224044 | 0 | 0 | proteasome |
| **Rpn8** | 1.889092017 | 0 | 0 | 0 | 0 | proteasome |
| **RPN9** | 1.584399756 | 0.298953662 | 0.273224044 | 0 | 4.62962963 | proteasome |
| **RPT4** | 1.462522852 | 3.886397608 | 4.918032787 | 4.895104895 | 28.7037037 | proteasome |
| **rpn6** | 1.157830591 | 0.298953662 | 2.459016393 | 0.932400932 | 9.259259259 | proteasome |
| **PRE8** | 1.096892139 | 0.298953662 | 2.18579235 | 1.631701632 | 14.81481481 | proteasome |
| **RPN1** | 1.035953687 | 2.69058296 | 6.284153005 | 7.692307692 | 24.07407407 | proteasome |
| **rpn11** | 0.914076782 | 0 | 0.819672131 | 0 | 5.555555556 | proteasome |
| **Ecm29** | 0.731261426 | 0 | 0 | 0 | 0 | proteasome |
| **Rpt1** | 0.548446069 | 4.783258595 | 5.737704918 | 0 | 1.851851852 | proteasome |
| **PRE3** | 0.548446069 | 1.046337818 | 3.005464481 | 1.165501166 | 6.481481481 | proteasome |
| **Rpn3** | 0.426569165 | 0 | 0 | 1.165501166 | 5.555555556 | proteasome |
| **RPN10** | 0.426569165 | 1.943198804 | 1.639344262 | 1.631701632 | 5.555555556 | proteasome |
| **rpn2** | 0.243753809 | 6.427503737 | 5.191256831 | 4.895104895 | 14.81481481 | proteasome |
| **PRE10** | 0.243753809 | 1.046337818 | 1.366120219 | 1.631701632 | 6.481481481 | proteasome |
| **rpn7** | 0.060938452 | 0.448430493 | 0 | 0 | 0 | proteasome |
| | 44.78976234 | 41.10612855 | 55.73770492 | 40.55944056 | 187.037037 | |
| | 23 | 17 | 19 | 13 | 18 | |
| **rrp4** | 0.731261426 | 0.298953662 | 0 | 0 | 0 | RNA decay |
| **rrp6** | 0.426569165 | 0.597907324 | 1.639344262 | 1.631701632 | 12.96296296 | RNA decay |
| **Sup35** | 1.279707495 | 0 | 0.546448087 | 0.233100233 | 0.925925926 | RNA decay |
| **LSM2** | 0.121876904 | 1.195814649 | 1.092896175 | 0.932400932 | 9.259259259 | RNA decay (U6) sno +rRNA processing |
| **LSM6** | 0.060938452 | 1.046337818 | 1.092896175 | 0 | 6.481481481 | RNA decay (U6) sno +rRNA processing |
| | 2.620353443 | 3.139013453 | 4.371584699 | 2.797202797 | 29.62962963 | |
| | 5 | 4 | 4 | 3 | 4 | |
| **CDC14** | 0 | 0 | 1.639344262 | 1.398601399 | 5.555555556 | rDNA organization |
| **NET1** | 0 | 0 | 1.639344262 | 1.398601399 | 15.74074074 | pol1 |
| **POL1** | 0.231261426 | 0.597907324 | 1.092896175 | 0.699300699 | 0 | pol1 |
| | 0.231261426 | 0.597907324 | 4.371584699 | 3.496503497 | 21.2962963 | |
| | 1 | 1 | 3 | 3 | 2 | |
| **TOR1** | 0.182815356 | 0.448430493 | 5.737704918 | 4.195804196 | 46.2962963 | Rbigenesis |
| **Nop1** | 0 | 3.437967115 | 7.103825137 | 5.827505828 | 21.2962963 | R biogenesis L |
| **Rpl8A** | 0 | 4.633781764 | 3.551912568 | 3.72960373 | 0 | R biogenesis L |
| **Drs1** | 0 | 0.149476831 | 0 | 0 | 1.851851852 | R biogenesis L |
| **Utp13** | 0 | 0 | 0 | 0 | 1.851851852 | R biogenesis L |
| **RTK1** | 0 | 0.298953662 | 1.639344262 | 0.932400932 | 11.11111111 | Rbigenesis |
| **UTP21** | 0 | 0.298953662 | 0.546448087 | 0 | 0 | R biogenesis L |
| **RIX7** | 0 | 0.597907324 | 2.459016393 | 0 | 0.925925926 | pre ribosomal export |
| **RRP3** | 3.900060938 | 0 | 0 | 0 | 0 | Rbigenesis |
| **KRE33** | 0.548446069 | 1.34529148 | 1.092896175 | 0.699300699 | 1.851851852 | R biogenesis S |
| **TSR1** | 0.365630713 | 0.298953662 | 0.273224044 | 0.466200466 | 11.11111111 | R biogenesis S |
| **DBP9** | 0.182815356 | 0 | 0 | 0 | 0 | Rbigenesis |
| **UTP20** | 0.182815356 | 0 | 1.366120219 | 0.466200466 | 0 | R biogenesis L |
| **ESF1** | 0 | 1.195814649 | 0 | 0.466200466 | 2.777777778 | R biogenesis L |
| **RPL30** | 0.670322974 | 1.943198804 | 4.918032787 | 3.03030303 | 20.37037037 | R biogenesis L |
| **ACL4** | 0.243753809 | 0 | 0 | 0 | 0 | R biogenesis L |
| **RPL1B** | 0 | 0 | 0.546448087 | 1.165501166 | 1.851851852 | R biogenesis L |
| **Utp23** | 0 | 2.69058296 | 6.284153005 | 3.962703963 | 19.44444444 | R biogenesis S |
| | 6.276660573 | 17.33931241 | 35.51912568 | 24.94172494 | 140.7407407 | |
| | 8 | 12 | 12 | 11 | 12 | |
| **RPL18A** | 0 | 0 | 0 | 0 | 1.851851852 | 60S protein |
| **RPL23B** | 0 | 0.747384155 | 1.092896175 | 1.165501166 | 4.62962963 | 60S protein |
| **L19** | 2.742230347 | 2.242152466 | 14.48087432 | 0 | 0 | 60S protein |
| **RPL12A** | 1.706276661 | 2.989536622 | 3.005464481 | 3.263403263 | 3.703703704 | 60S protein |
| **RPL5** | 1.340645948 | 4.484304933 | 5.191256831 | 4.662004662 | 5.555555556 | 60S protein |
| **RPL23B** | 1.035953687 | 1.34529148 | 0.819672131 | 0.932400932 | 5.555555556 | 60S protein |
| **RPL25** | 0.975015235 | 0 | 0 | 0 | 0 | 60S protein |

| | | | | | | |
|---|---|---|---|---|---|---|
| **RPP0** | 0.914076782 | 0 | 0 | 0 | 0 | 60S protein |
| **RPL30** | 0.670322974 | 1.943198804 | 4.918032787 | 3.03030303 | 20.37037037 | 60S protein |
| **RPL33A** | 0.426569165 | 2.69058296 | 3.825136612 | 4.895104895 | 7.407407407 | 60S protein |
| **RPP2A** | 0.182815356 | 0.298953662 | 0 | 0 | 8.333333333 | 60S protein |
| **RPL5** | 0.060938452 | 5.381165919 | 2.18579235 | 0.466200466 | 0 | 60S protein |
| **RPL18** | 0 | 0 | 0 | 0 | 1.851851852 | 60S protein |
| **RPS26A** | 6.154783668 | 1.046337818 | 0 | 0 | 0 | 40S protein |
| **RPS15** | 1.096892139 | 3.288490284 | 2.18579235 | 4.428904429 | 0 | 40S protein |
| **RPS10B** | 0.792199878 | 3.886397608 | 0 | 10.72261072 | 0 | 40S protein |
| **ASC1** | 0.609384522 | 0 | 0 | 0 | 0 | 40S protein |
| **RPS7A** | 0.365630713 | 0 | 0 | 0 | 0 | 40S protein |
| **RPS9B** | 0.243753809 | 0.896860987 | 0.819672131 | 6.526806527 | 15.74074074 | 40S protein |
| **RPS0B** | 0.182815356 | 0.298953662 | 0.819672131 | 0 | 0 | 40S protein |
| **RPS7B** | 0 | 0 | 0.273224044 | 0 | 0 | 40S protein |
| **Frs1** | 0 | 0 | 0 | 0.699300699 | 3.703703704 | tRNAprocessing |
| **RET1** | 0 | 2.69058296 | 8.196721311 | 5.594405594 | 36.11111111 | pol III |
| **Sap185** | 0 | 15.09715994 | 2.732240437 | 0.233100233 | 1.851851852 | tRNAprocessing |
| **RPC53** | 0.609384522 | 0.597907324 | 1.092896175 | 0.699300699 | 2.777777778 | pol III |
| **MES1** | 0 | 0.298953662 | 0.273224044 | 0 | 3.703703704 | tRNA nucelar export |
| **Los1** | 0.670322974 | 0 | 1.092896175 | 0.466200466 | 0.925925926 | Nuclear pore protein; involved in nuclear export of pre-tRNA |
| **Pol32** | 0.182815356 | 0 | 0 | 0 | 3.703703704 | pol III |
| **LSM2** | 0.121876904 | 1.195814649 | 1.092896175 | 0.932400932 | 9.259259259 | tRNAprocessing |
| | 1.584399756 | 19.88041854 | 14.48087432 | 8.624708625 | 62.03703704 | |
| | 4 | 5 | 6 | 6 | 8 | |
| **Bcd1** | 8.653260207 | 0 | 0 | 0 | 0 | Sno Processing |
| **UTP14** | 0 | 0.597907324 | 1.366120219 | 1.398601399 | 0 | Sno Processing |
